# Detection of Somatic Point Mutations Directly from Spatial Transcriptomics Enables in vivo Spatiotemporal Lineage Tracing

**DOI:** 10.64898/2026.02.04.703493

**Authors:** Zhirui Yang, Mengdie Yao, Qing Yang, Yiheng Du, Jinhong Lu, Xing Wu, Jinran Lin, Zhuoyu Qian, Song Hu, Yonghe Xia, Huan Liu, Qiji Zhou, Xiaoyu Ma, Yan Luo, Wenxuan Fan, Weike Pei, Yuhao Xia, Xiuyan Yu, Jing Luan, Qiao’an Zhang, Yue Zhang, Qin Wang, Jianyang Zeng, Yue Zhang, Wenyu Wu, Yanmei Dou

**Affiliations:** School of Life Sciences, Westlake University, Hangzhou, Zhejiang, 310024, China; Westlake Laboratory of Life Sciences and Biomedicine, Hangzhou, Zhejiang, 310024, China; College of Life Sciences, Zhejiang University, Hangzhou, Zhejiang, China; Fudan University, Shanghai, 200433, China; Department of Dermatology, Huashan Hospital Fudan University, Shanghai, China; Department of Dermatology, Jing’an District Central Hospital, Shanghai, China; Department/Institute of Neurobiology, School of Basic Medical Sciences, Xi’an Jiaotong University Health Science Center, Xi’an 710061, China; School of Engineering, Westlake University, Hangzhou, Zhejiang, China; Department of Breast Surgery, Second Affiliated Hospital, Zhejiang University, Hangzhou, Zhejiang, China

**Author notes:** Correspondence (W.W.), (Y.Dou). These authors contributed equally (Zhirui Yang, Mengdie Yao, Qing Yang, Yiheng Du, Jinhong Lu).

## Abstract

Spatial transcriptomics reveals tissue organization but lacks in vivo lineage-tracing methods applicable to humans. We introduce SpaceTracer, a computational framework that accurately detects somatic single-nucleotide variants (SNVs) directly from spatial transcriptomics data. By leveraging naturally occurring somatic SNVs, SpaceTracer reconstructs cellular phylogenies within native tissue architecture, enabling the mapping of lineage spread, migration, lineage-coupled expression changes and lineage-aware local interactions. Applied to human cutaneous squamous cell carcinoma, it traced tumor initiation and progressions, uncovered widespread pre-invasive migration of dedifferentiated epithelial cells and characterized mutant B cells migrating from tertiary lymphoid structures (TLS) into the tumor boundary. The framework also reconstructed developmental lineages across multiple tissues and identified tissue-resident mutant immune cells. SpaceTracer thus provides a perturbation-free platform for high-resolution spatiotemporal lineage tracing, offering a transformative tool for elucidating complex biological systems—especially tumor-immune ecosystems—with direct implications for advancing cancer immunotherapy.

## Introduction

Somatic mutations play a pivotal role in many biological processes, such as cancer progression, aging, and disease evolution, and could serve as in vivo markers for lineage tracing^1,2^. Traditionally, the detection of somatic mutations has relied on high-coverage genomic sequencing data. The advent of spatial sequencing data provides a powerful tool to investigate the intricate relationships between somatic mutations, cellular function, genetic variation, and tissue architecture.

Although a few methodologies exist to call somatic copy number variations (CNVs) directly from spatial sequencing data^3,4^, these methods face significant limitations when decoding spatial clonal architecture. First, CNVs often lack the resolution needed to capture fine-grained clonal dynamics, particularly in cancer types and benign tumor samples where CNV abundance is low^5,6^. Second, current CNV detection methods based on transcriptomics data typically assume that non-cancer cells have diploid genomes, limiting their ability to delineate clonal structures in normal tissues^3,7–9^. Lastly, detection of CNVs can be highly influenced by gene expression patterns and the varying numbers of cells within different spatial spots. In contrast, somatic SNVs are much more prevalent in both tumor and normal samples and typically occur as binary events (non-mutant or mutant), making them less affected by these issues. While somatic SNVs are powerful markers for high-resolution clonal reconstruction, their potential in spatial transcriptomics remains untapped. Existing methodologies—whether designed to detect somatic SNVs from single cell RNAseq data^10–12^, or methods repurposed from DNA-based analysis^13^ lack the precision and sensitivity needed to reliably identify these variants in spatial data, fundamentally limiting accurate lineage tracing.

Spatial sequencing data typically encompass a relatively small portion of the genome, and this inherent sparsity has led to skepticism regarding its ability to reveal detailed mutational landscapes^14^. Furthermore, the noise in RNAseq data complicates the identification of subtle genomic alterations, such as somatic SNVs, making the development of a reliable detection algorithm challenging^15^. However, “deep and sparse” sequencing may prove more effective than “shallow and broad” sequencing, particularly for somatic mutation detection and single-cell lineage tracing. Although spatial transcriptomics capture a smaller fraction of the genome, the read coverage is typically very deep within the limited regions of a small number of cells (Fig. 1a), allowing the detection of somatic SNVs in small clonal populations and enabling precise lineage reconstruction. In contrast, bulk WGS sequencing provides more even coverage across the genome, which may help identify somatic mutations across broader genomic regions. However, this approach substantially reduces the ability to detect mutations in smaller clones and to reconstruct phylogenetic trees (Fig. 1a). Furthermore, the spatial organization of cells offers a potential advantage in mitigating noise within the data (Fig. 1, b-c), providing a new avenue for mutation detection.

**Fig. 1.**
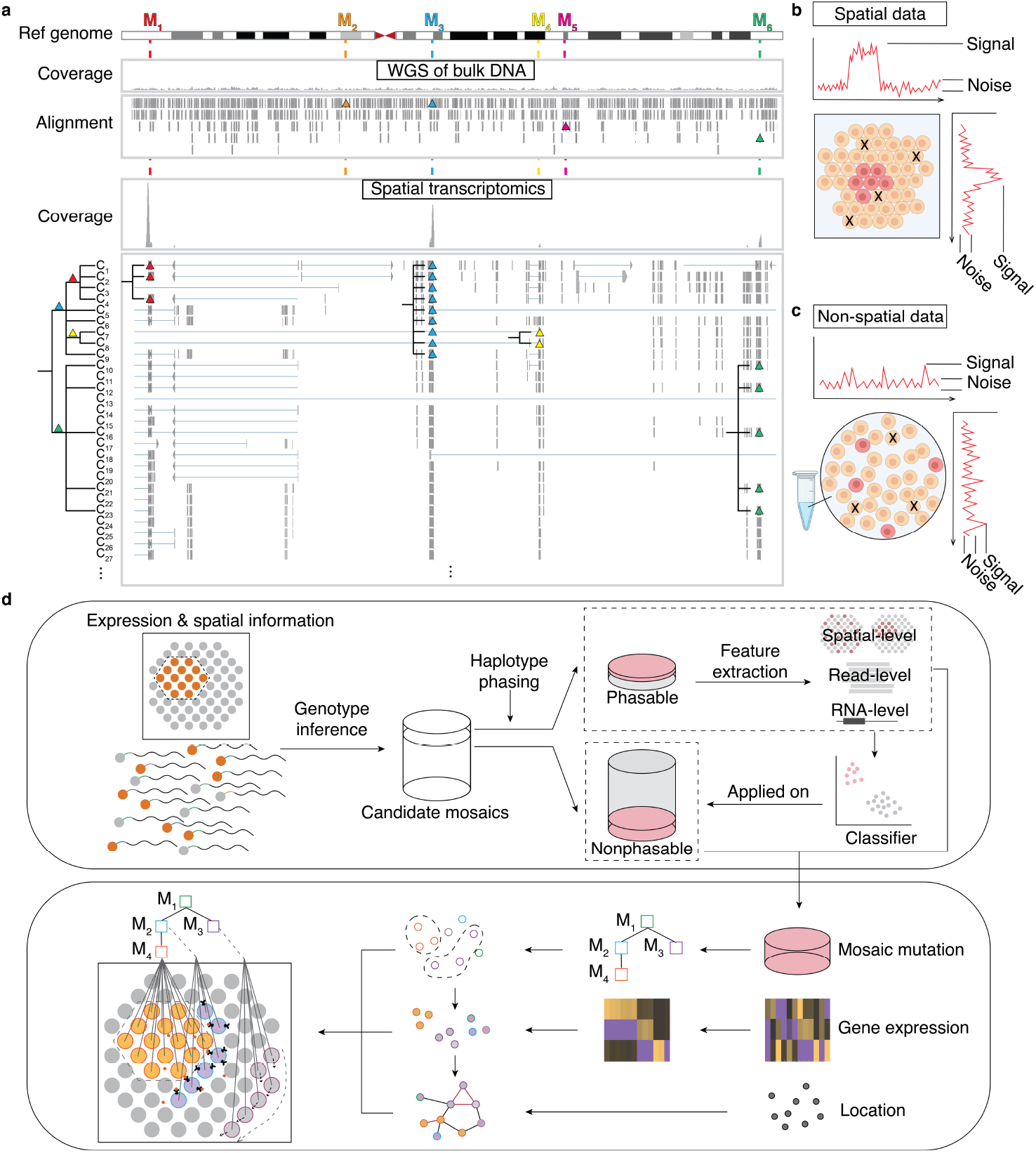
Overview of SpaceTracer. (**a**) Comparison of ‘deep and sparse’ sequencing with ‘shallow and broad’ sequencing for somatic mutation detection and single-cell lineage tracing. Spatial transcriptomics capture a smaller fraction of the genome but provide deep read coverage in limited regions of a small number of cells, enabling the detection of somatic SNVs in small clonal populations and facilitating precise lineage reconstruction. In contrast, bulk WGS sequencing offers more even coverage across the genome from a large number of cells, which aids in identifying somatic mutations in broader genomic regions, but significantly reduces the ability to detect mutations in smaller clones and to reconstruct phylogenetic trees. The triangles indicate somatic mutations. (**b-c**) The spatial arrangement of cells offers an advantage in reducing noise, as adjacent cells are more likely to share common mutations due to their lineage relationships. By utilizing spatial proximity, the local signal-to-noise ratio can be enhanced, providing a more robust platform for mutation detection. (**d**) Hierarchical Bayesian inference was applied to screen for candidate somatic mutations from raw sequencing data. Haplotype phasing, along with spatial-, read-, and site-level features, was used to train a model for classifying high-confidence somatic variants. These variants were subsequently used for spatial lineage tracing, enabling spatial-temporal lineage reconstruction and the integration of lineage, spatial, and gene expression data into a unified framework. This approach facilitates the study of complex biological processes, such as development, cell migration, and cell-cell communication.

Based on this rationale, we developed SpaceTracer, a novel algorithm capable of accurately detecting somatic SNVs directly from spatial transcriptomics data. Across multiple benchmarks, our approach consistently achieved accuracy rate of 80% or higher. Using spatial transcriptomics data from both tumor and normal samples, SpaceTracer successfully identified a couple to hundreds of somatic SNVs per sample, which enables lineage tracing in space. These findings enable us to decode a tumor’s cell-of-origin, trace lineage-aware cellular migrations, and map the dynamic evolution and tumor-immune interactions within tumor ecosystem. Furthermore, they allow us to explore key developmental and communication processes in both healthy and diseased human tissues. This work demonstrates the potential of spatial sequencing data as a rich source of mutational data, revealing insights previously obscured by technical limitations. SpaceTracer is open-source, licensed under the MIT License, and publicly accessible on GitHub at https://github.com/douymLab/SpaceTracer. The SpaceTracer source code is being prepared for public release. It will be available on GitHub by March 1, 2026, or upon the manuscript’s formal acceptance, whichever comes first.

## Result

### Framework of SpaceTracer

SpaceTracer is a robust algorithm that combines advanced statistical frameworks, haplotype phasing, and machine learning techniques to detect somatic SNVs directly from spatial sequencing data (Fig. 1d, Extended Data Fig. 1a, Methods). The basis of SpaceTracer’s methodology involves a statistically principled hierarchical Bayesian network, which is employed to scan for candidate loci within the spatial datasets (Extended Data Fig. 1b, Methods). This approach allows us to systematically identify sites of interest that may harbor mutations.

To enhance the accuracy of mutation detection, we designed and extracted nearly a hundred statistical, proper-normalized read-level, spatial-distribution-related features and RNA-specific metrics from candidate loci, specifically designed to distinguish real somatic mutations from artifacts (Extended Data Fig. 1c-d, Supplementary Table 1). These features were carefully chosen for their ability to provide reliable metrics on the confidence of candidate loci. A machine-learning model was built with these features to predict somatic mutations based on a rigorously curated training set with orthogonal validations (Methods). To further distinguish these predictions from artifacts that mimic real mutations—such as chemical modifications and RNA-editing events—we applied an artifact mutation signature analysis. This enabled the systematic filtering of lysis errors and other technical noise from the RNA data (Extended Data Fig. 2, Methods).

### Benchmark of SpaceTracer

To achieve an unbiased evaluation of mutation detection, we established a benchmark dataset through a dual-strategy approach. First, we generated paired spatial transcriptomic and deep-coverage (200×) whole-genome sequencing (WGS) data from adjacent tissue sections of a basal cell carcinoma (BCC) sample (Fig. 2a, Methods). This was supplemented by curating six samples from three independent, publicly available spatial transcriptomics datasets^16–18^ (Supplementary Table 2), selected specifically for the availability of sequencing data from neighboring sections—including single-cell RNA sequencing (scRNA-seq) or spatial profiles (Methods). Model performance was then evaluated using a leave-one-donor-out cross-validation strategy (Methods). The performance of SpaceTracer was compared with three established methods, including (1) Monopogen^10^ and (2) Scomatic^11^, both originally developed for detecting somatic SNVs in single-cell RNA-seq data and adapted here for spatial transcriptomics (Methods); and (3) SpatialSNV^13^, which calls somatic mutations from spatial data using Mutect2^19^. It should be noted that, as these methods were not optimized for somatic mutation detection in spatial transcriptomics data, additional filters were applied to remove low-quality variant calls (Methods).

**Fig. 2.**
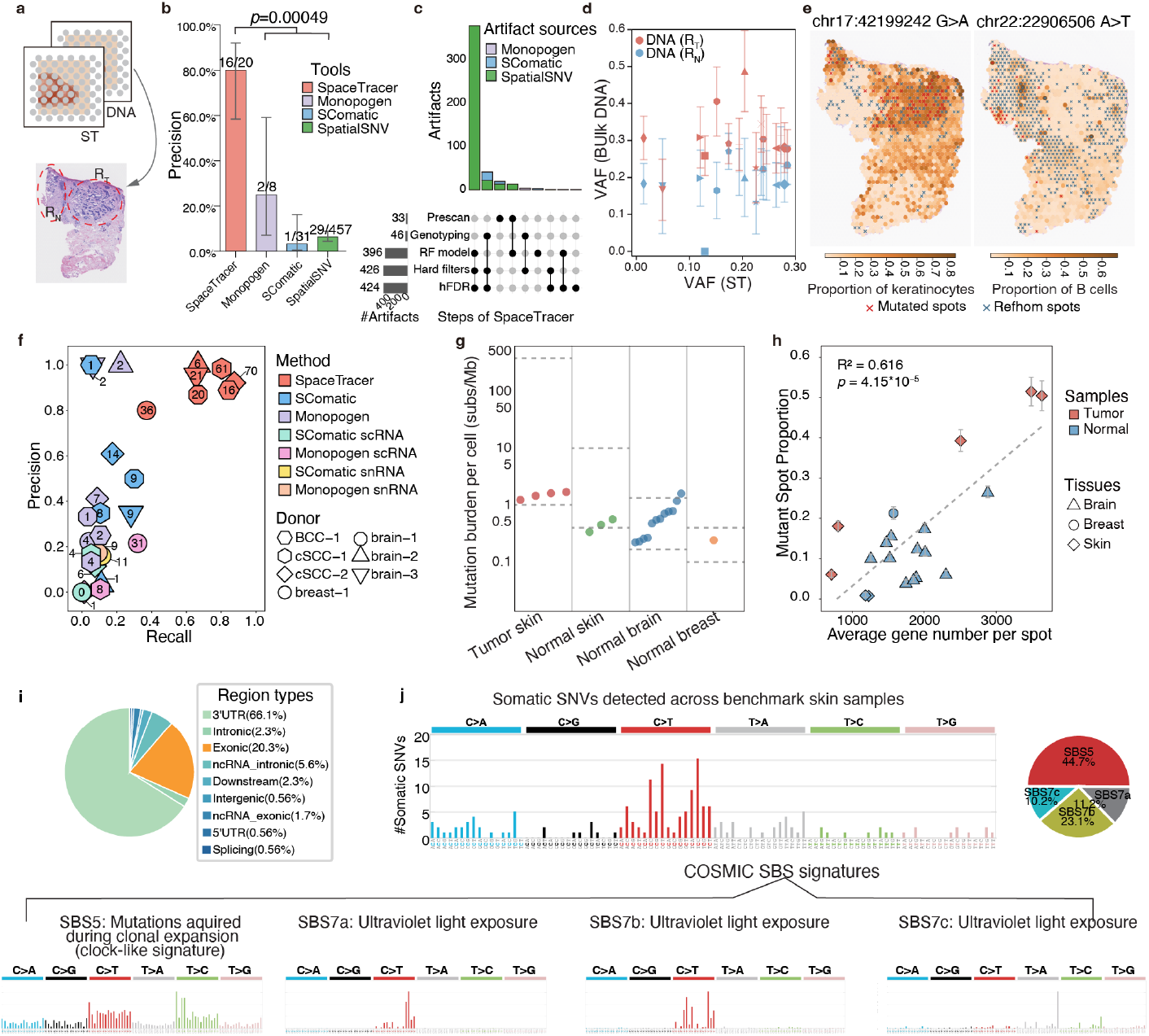
Benchmarking of mutation detection. (**a**) Schematic illustrating the processing of a Basal Cell Carcinoma (BCC) sample for benchmarking. One tissue slice was used for spatial transcriptomics, while a consecutive slice underwent deep WGS sequencing. Regions of interest (R_T_: tumor; R_N_: mostly normal), identified from the H&E staining, were micro-dissected and sequenced separately to 200× coverage. (**b**) SpaceTracer demonstrated substantially higher accuracy than other methods. Error bars represent 95% confidence intervals derived from binomial sampling. P-value was calculated with one-tailed t-test. Values above bars indicate performance metrics, which represents validated calls out of total calls made by the tool. (**c**) Artifact variants identified by other software tools were effectively excluded by the comprehensive framework of SpaceTracer, including the steps of pre-scan, Bayes genotyping, mutational signature deconvolution and a series of hard filters. (**d**) Validation of somatic SNVs called by SpaceTracer. Variant allele frequencies (VAFs) measured by deep DNA sequencing are shown for the Rt region (red) and the Rn region (blue). Error bars represent 95% confidence intervals of VAFs derived from binomial sampling. (**e**) Spatial distribution of two somatic SNVs uniquely detected by SpaceTracer across multiple datasets. Red Xs mark mutated cells, and blue circles mark reference homozygous cells with non-zero coverage. (**f**) Superior precision and recall of SpaceTracer. In evaluations using our benchmark data and independent public datasets with sequencing from neighboring tissues, SpaceTracer demonstrated optimal performance in both precision and recall. The set of real mutations from public datasets was defined as the union of cross-validated somatic SNVs identified by SpaceTracer, Monopogen, and SComatic (see Methods for more details). The number of real mutations detected by each tool per sample is indicated within its corresponding shape. (**g**) Mutation rates across tissues are consistent with published ranges. Somatic mutations were identified in various tissue samples, and the per-cell mutation rates for all samples fell within the range (horizontal dashed lines) established by prior DNA sequencing studies. See Methods for details. (**h**) Greater sequencing depth enhances the detection of mutant cells. Mutations used to classify a spot as mutant were identified genome-wide by SpaceTracer. Error bars represent 95% CIs derived from binomial sampling. (**i**) Functional annotation of somatic SNVs detected across benchmark skin samples. (**j**) Mutation signatures extracted from the benchmark skin samples align with known biological processes and published mutation signatures from DNA sequencing.

Accurate and sensitive somatic SNV detection is fundamental to high-resolution lineage tracing. Benchmarked against deep-coverage DNA sequencing data from adjacent tissue sections, SpaceTracer achieved significantly higher accuracy than competing methods (p=0.00049, two-tailed t test, Fig. 2b–e, Extended Data Fig. 3a-d, Supplementary Table 3, Methods). Of note, we trained two separate models, one incorporating spatial distribution features and another omitting them. The model without spatial features achieved comparable precision in mutation calling (Extended Data Figs. 1d & 3c). This demonstrates the broader applicability of SpaceTracer: it remains effective even in datasets where clonal populations lack clear spatial organization, such as in contexts involving extensive cell migration. On six independent, publicly available datasets spanning tumor and normal samples, SpaceTracer consistently outperformed other callers in accuracy (overall≥80%, p=3.9*10^-9^, one-tailed t-test, Extended Data Fig. 4e) while also exhibiting top-tier sensitivity (p=1.45*10^-4^, one-tailed t-test, Extended Data Fig. 3f-h, Supplementary Table 4, Methods).

Across these benchmark datasets, SpaceTracer identified several to dozens of somatic SNVs per tissue slice (Fig. 2g-h), with ∼10-20% in exonic regions (Fig. 2i, Extended Data Fig. 4). The per-cell mutation rates estimated from these calls align closely with published rates from DNA sequencing^20–23^ (Fig. 2g; Methods). Moreover, the mutational signatures extracted from the spatial data matched known DNA-based signatures (Fig. 2j, Extended Data Fig. 4, Methods). Together, these results validated the robustness and accuracy of SpaceTracer’s mutation detection.

To benchmark phylogenies inferred from spatial somatic SNVs, we established two ground-truth datasets by projecting cells from public CRISPR-barcoded LUAD scRNA-seq datasets^24^ onto a spatial layout (Fig. 3, Methods). Phylogenetic trees were then reconstructed from the resulting mutation profiles using PhyloSOLID^25^, a software tool we developed for robust phylogeny reconstruction that tolerates the inherent sparsity and noise of single-cell data. In both benchmarks, our pipeline achieved accurate lineage reconstruction, which was highly consistent with the CRISPR barcode trees (Fig. 3, Extended Data Fig. 5).

**Fig. 3.**
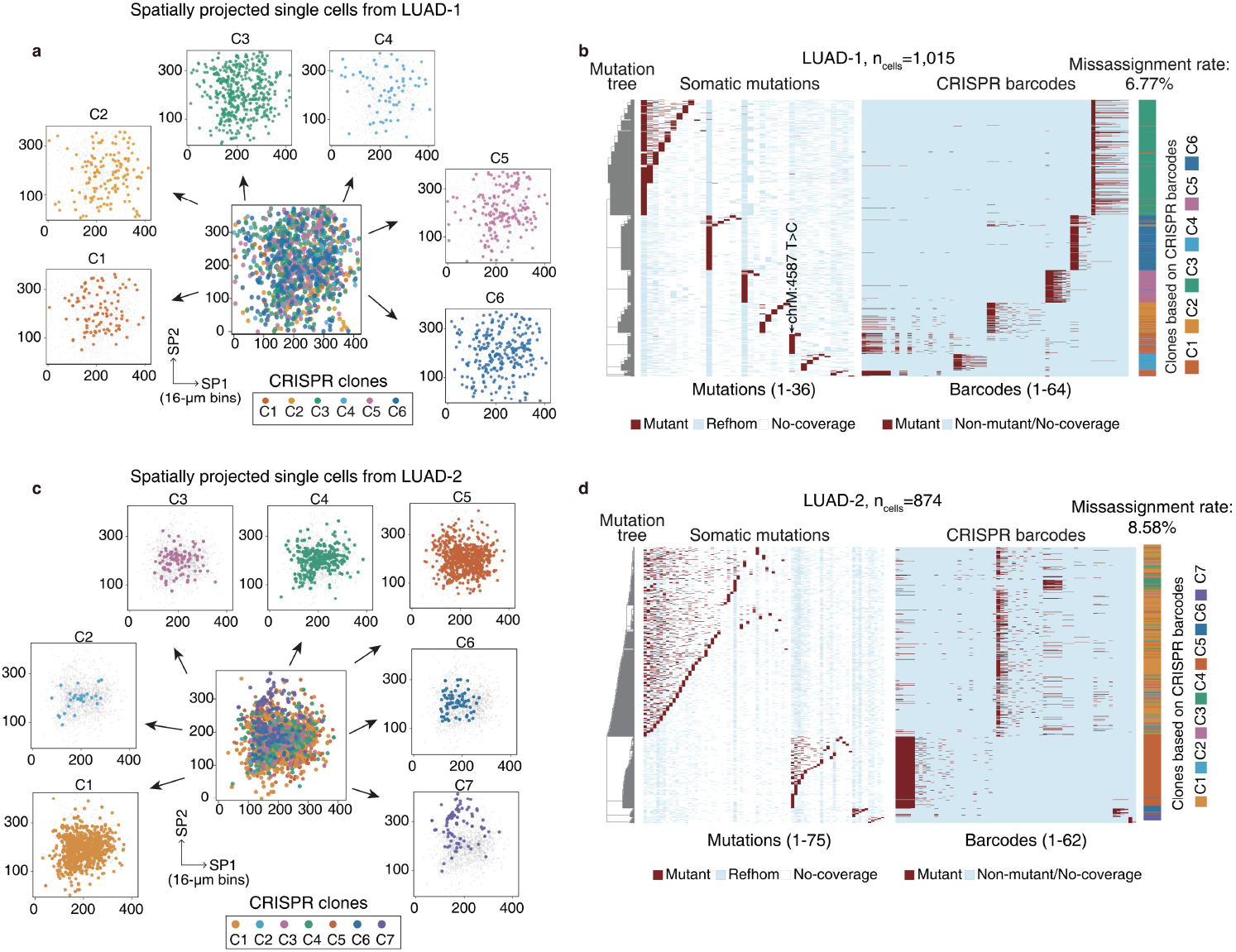
Benchmarking of phylogeny reconstruction. (**a**) Spatial projection of CRISPR-barcoded single cells from LUAD-1. Single-cell RNA-seq data from cells with unique CRISPR barcodes were projected onto a spatial layout. Colors correspond to distinct barcoded clones. See Methods for details. (**b**) Phylogenetic concordance between somatic SNV derived and ground-truth lineages of LUAD-1. The phylogeny reconstructed from spatially projected single-cell data (left) shows high concordance with the ground-truth CRISPR barcode clones (right). In the heatmaps, rows represent single cells and columns represent individual somatic or barcode mutations. (**c**) Spatial projection of CRISPR-barcoded single cells from LUAD-2. (**d**) Phylogenetic concordance between somatic SNV derived and ground-truth lineages of LUAD-2.

### Decoding tumor initiation in cutaneous squamous-cell carcinoma (cSCC)

To demonstrate the ability of SpaceTracer in decoding tumor lineages at spatial resolution, we selected cSCC as a foundational benchmark. The characteristic histological progression of cSCC allows a single tissue section to encompass the complete tumorigenic continuum providing an ideal model for spatially resolved analysis. From a published cSCC dataset^18^, our analysis detected nearly one hundred somatic SNVs across two slices of one individual sample cSCC-1, of which ∼20% were exonic mutations (Supplementary Table 5). To characterize the cellular composition of each spot, we performed expression-based deconvolution (Supplementary Table 6, Methods), distinguishing epithelial cells, keratinocytes and immune cells. Epithelial cells and keratinocytes were further distinguished into tumor cells and normal cells by using tumor-specific keratinocyte (TSK) score (Methods). Using somatic SNVs detected, we reconstructed a hierarchical phylogenetic tree (Methods, Fig. 4a). The tree clearly delineated clusters corresponding to tumor, normal, and immune-associated spots, revealing distinct lineage relationships among these populations.

**Fig. 4.**
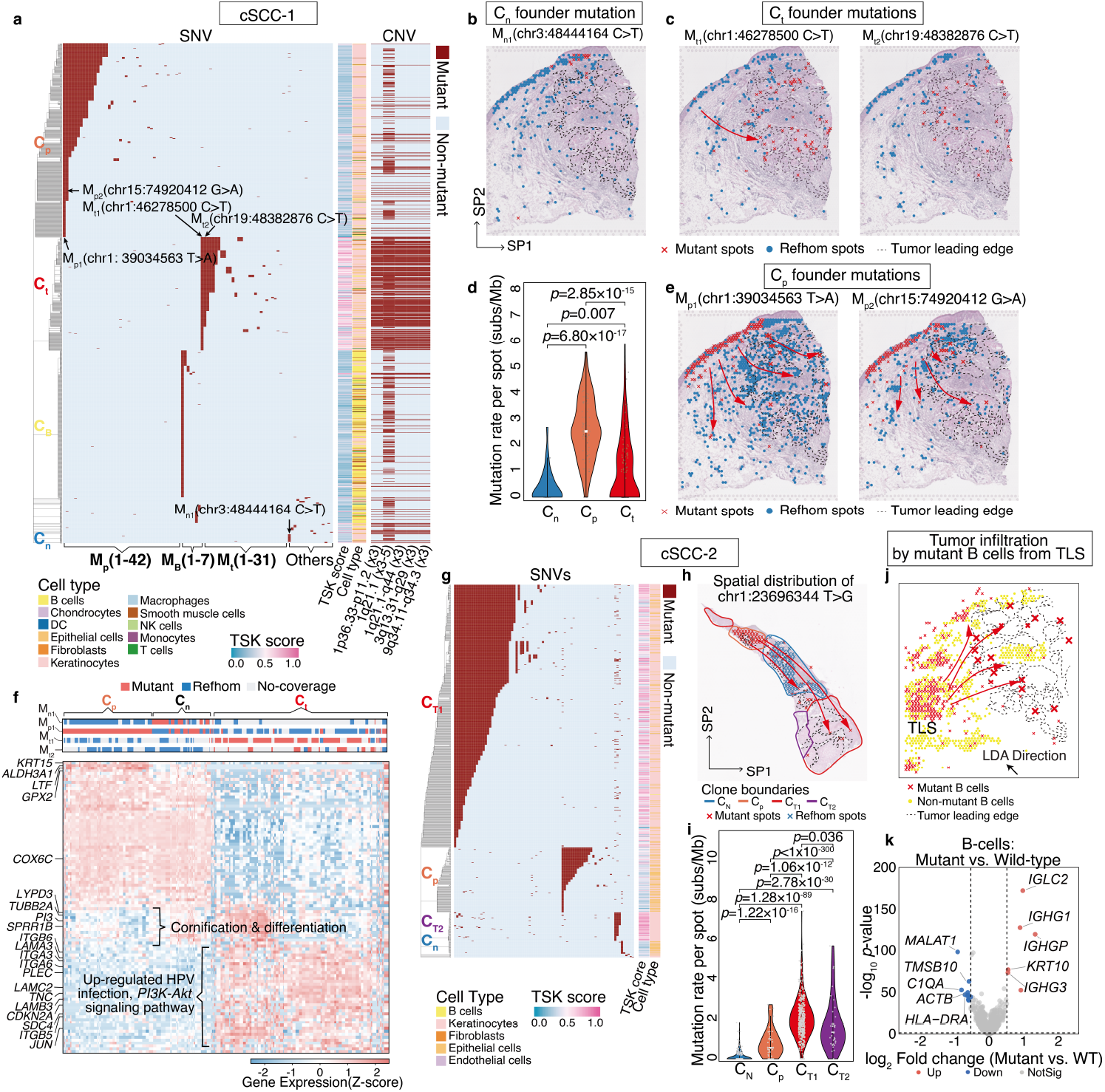
Key biological insights revealed by SpaceTracer in cSCC. (**a**) Phylogeny reconstructed using somatic SNVs from two slices of cSCC-1 reveals the cellular composition and lineage divergence within the tumor. Tumor keratinocytes, normal epithelial cells, and immune cells are clustered into three distinct clones. The heatmap displays the mutational profile for each spatially resolved spot (rows) across the identified somatic SNVs (columns). Spots that are phylogenetically imputed as mutant, as well as those inferred from a mutant -> reference-homozygous pattern, are both annotated as “mutant”. This uniform annotation is designed to accommodate the potential co-capture of cells from multiple lineages within a single spot. The profile of CNVs is shown on the right. (**b**) Spatial distribution of the normal epithelial clone Cn identified in panel (**a**). Spots are annotated based on the Cn-unique founder mutation chr3:48444164 C>T. Red Xs mark mutant spots; blue circles mark reference-homozygous (refhom) spots. (**c**) Spatial distribution of the tumor epithelial clone C_t_, traced with two C_t_-unique founder mutations. (**d**) Per-cell mutation rate across epithelial clones of cSCC-1. P-values were calculated with two-tailed t tests. (**e**) Differentially expressed genes and pathways across clones. The heatmap displays gene expression (rows) for each cell (columns). Cells were clustered based on the Euclidean distance of the top 100 differentially expressed genes. See Methods for details. (**f**) Spatial distribution of the migrating non-tumor epithelial clone C_p_, traced with two C_p_-unique founder mutations. The mutations M_t1_, Mt_2_, M_p_, M_n_ correspond to the clone-specific founder mutations shown in panel **a**. (**g**) Phylogeny reconstructed using somatic SNVs from two slices of an independent sample, cSCC-2, reveals polyclonal tumor of origin. (**h**) A migrating non-tumor epithelial clone C_p_ from cSCC-2, traced with a C_p_-specific founder mutation. (**i**) Per-cell mutation rate across epithelial clones of cSCC-1. P-values were calculated with two-tailed t tests. (**j**) Mutant and non-mutant B cells exhibit distinct spatial distributions (p = 5.7 × 10^−20^, Kolmogorov-Smirnov test along the LDA direction). The Linear Discriminant Analysis (LDA) direction represents the spatial axis of greatest difference in cell density between the two groups, as visualized in the original panel. Mutant B cells (red Xs) are shown emigrating from a TLS and infiltrating the tumor region (dashed lines), whereas non-mutant B cells (yellow, identified by deconvolution as the major cell type within the spot) are predominantly retained in the peritumoral space. The large red Xs represent mutant B cells infiltrating the tumor boundary, while the small red Xs denote hypermutant B cells located outside of tumor region. (**k**) Analysis of differential gene expression of mutant B cells versus non-mutant B cells.

Since CNVs account for a significant portion of the genetic basis of cancer^5^, we also inferred phylogenetic lineages from CNVs identified in the spatial data using calicoST^4^ and inferCNV^26^ (Fig. 4a, Extended Data Fig. 6, Methods). Although the CNV-based tree broadly separated tumor from normal clones—with minor discrepancies attributable to the multi-cellular composition of Visium spots (Extended Data Fig. 7)—its limited informative CNVs produced a low-resolution phylogeny. In this tree, most non-tumor spots shared one identical CNV profile, and most tumor spots shared another. In contrast, the somatic SNVs detected by SpaceTracer enabled the delineation of refined tumor subclones and revealed distinct clonal structures among non-tumor cells.

Investigating tumor initiation is crucial for understanding early carcinogenesis and remains one of the most critical challenges in cancer research^27^. Unlocking this process is akin to identifying the “first domino” in the cascade of cancer progression, with profound clinical implications. To explore the spatiotemporal dynamics of clonal initiation, we focused on three genetically distinct epithelial clones: C_n_, C_p_, and C_t_ (Fig. 4a). As shown, cells of the C_n_ clone (marked by C_n_-specific founder mutation M_n1_) are largely confined to the epithelium (Fig. 4b). In contrast, tumor-forming cells of the C_t_ clone (marked by a C_t_-specific founder mutation M_t1_) likely migrated from the epithelium into the dermis, subsequently expanding to form the bulk of the tumor (Fig. 4c). Temporal analysis utilizing phylogeny indicates C_t_ cells likely migrated from the epithelial region to the tumor region, as most tumor spots harbor the M_t_ mutation and have accumulated additional sub-clonal mutations (Fig. 4a & c). Within these tumor regions, the mutation rate per cell is significantly elevated compared to normal epithelial cells of the C_n_ lineage (p=0.007, two-tailed t-test, Fig. 4d, Extended Data Fig. 8a-b, Methods). Furthermore, gene expression analysis reveals significant upregulation of HPV infection and *PI3K-Akt* signaling pathways within the Ct clone (Fig. 4e), both of which are classical pathways contributing to cSCC etiology^28^.

Intriguingly, C_p_—a lineage phylogenetically independent of both C_n_ and C_t_—was neither confined to the epithelium nor contributed to tumor formation. Despite this, C_p_-clone cells exhibited extensive migrations (Fig. 4f). Notably, their migratory trajectory spatially encompasses that of the tumor cells, indicating a non-tumorigenic cell migration event. Strikingly, the estimated per-cell mutation rate of Cp cells was even significantly higher than that of tumor cells (p=2.85*10^-15^, two-tailed t-test, Fig. 4e, Extended Data Fig. 8a-b, Methods). Gene expression analysis revealed that C_p_ cells have undergone significant down-regulation of cornification and cell differentiation processes (Fig. 4e, Methods).

Notably, the extensive migration of non-tumor epithelial cells was not an isolated event, as we recurrently observed the same phenomenon in an independent cSCC sample, cSCC-2 (Fig. 4g-h), suggesting it could be a pervasive feature of cSCC. Mirroring our findings in cSCC-1, the migrating non-tumor epithelial clone in cSCC-2 (C_p_) also exhibited a significantly elevated mutation rate per cell compared to other normal epithelial cells (p=1.22×10^−16^, two-tailed t-test; Fig. 4i). Moreover, lineage-coupled analysis (Extended Data Fig. 8c-d) revealed that the C_p_ clone demonstrated concomitant upregulation of key pro-migration genes: Dystonin (*DST*), 24-Dehydrocholesterol Reductase (*DHCR24*), and Immunoglobulin Superfamily Member 3 (*IGSF3*)^29–31^. Intriguingly, cSCC-2 is of polyclonal origin, with its two founding clones (C_T1_ and C_T2_) being phylogenetically distant, exhibiting significantly different mutation rate (p=0.036, two-tailed t test, Fig. 4i) and different transcriptional profiles (Extended Data Fig. 8e-f).

Collectively, beyond tracing tumor initiation and progression, our findings suggest that the widespread migration of non-tumor-forming epithelial cells in cSCC is linked to a distinct dedifferentiation event. This transformative state likely facilitates cell mobility, potentially aiding tumor invasion or fostering a pro-tumorigenic microenvironment. Crucially, this phenomenon was undetectable using gene expression data alone, underscoring the power of our somatic SNV-based methodology to reveal these migratory, low-abundance cell populations within each spatial spot.

### Spatial-resolved clonal lineages and immune interactions: decoding the ecosystems of tumor progression

Tumor subclones tend to expand at heterogeneous rates. To quantify these differences spatially, we computed the phylogenetic fitness score for each spatial transcriptomics spot based on the reconstructed phylogenetic tree using an established method^32^ (Methods). This score defines the subclonal growth rate. We observed that spots containing cells with higher phylogenetic fitness scores also showed elevated expression of both a gene-expression-based fitness signature (p=0.012, Spearman correlation analysis, Extended Data Fig. 9a) and a tumor epithelial–mesenchymal transition (EMT) score (p=0.026, Spearman correlation analysis, Extended Data Fig. 9b). This pattern suggests that rapidly expanding subclones adopt a more aggressive phenotypic state. Gene expression analysis revealed that tumor subclones with higher phylogenetic fitness scores significantly upregulate energy metabolism-related genes such as Mitochondrially Encoded NADH:Ubiquinone Oxidoreductase Core Subunit 5 (*MT-ND5*), supporting their proliferative and biosynthetic demands, while downregulating *CD151* to enhance metastatic and invasive potential (Extended Data Fig. 9c-d). Furthermore, expression levels of Leucine Rich Repeat Containing 41 (*LRRC41)*, DExD/H-Box Helicase 60 (*DDX60*), and PAT1 Homolog 1 (*PATL1)*—which correlate most strongly with phylogenetic fitness scores in linear regression analysis (Extended Data Fig. 9e-f, Methods) have known roles in mitosis and cancer pathogenesis^33–35^, implicating their potential involvement in the molecular mechanisms driving accelerated clonal expansion.

Cell–cell communication analysis of tumor keratinocytes revealed that rapidly proliferating tumor subclones (those with higher phylogenetic scores) markedly upregulate annexin A1 (*ANXA1*, Extended Data Fig. 9g-i, Methods). *ANXA1* is a key mediator of sterile inflammation and intercellular signaling known to promote macrophage polarization toward a pro-tumoral M2 phenotype. It has also been shown to suppress antigen-specific cytotoxic T cell responses and facilitate the induction of regulatory T lymphocytes^36^.

Furthermore, we identified seven somatic SNVs within B cell receptor (BCR) loci (Fig. 4j), 4/7 were located in BCR variable regions (*IGHV* and *IGLV*, Supplementary Table 7), suggesting somatic hypermutation in B cells. These mutated B cells exhibited upregulated expression of antibody genes (e.g., *IGLC2, IGHG1, IGHG*; Fig. 4k), consistent with an active plasma cell phenotype. Spatial analysis revealed a distinct distribution pattern: mutated B cells were highly enriched in peritumoral TLS and deeply infiltrated the malignant core, while non-mutated B cells were located outside of the tumor (Fig. 4j). Intriguingly, cell-cell communication analysis indicated an elevated expression of the *CD46* receptor on these mutated B cells alongside high Jagged-1 (*JAG1*) expression on tumor cells (Extended Data Fig. 9j). This ligand-receptor pair implies a potential tumor immune evasion strategy whereby tumors may co-opt the reverse *Notch* signaling pathway via *CD46-JAG1* interaction to suppress the function of tumor-infiltrating, antigen-experienced B cells^37^.

### Spatiotemporal decoding of tumor development and tissue-resident B cells

Beyond cSCC, we applied SpaceTracer to other tumor types, successfully detecting mutations and reconstructing spatiotemporal phylogenies. For instance, analysis of glioblastoma (Extended Data Fig. 10) delineates clear patterns of tumor initiation and progression, demonstrating the broad utility of our approach.

Tissue-resident immune cells are essential for maintaining tissue homeostasis^38^. To investigate immune cells residing in normal tissues and their associated mutations, we examined the prevalence of somatic mutations localized to immunoglobulin loci in sequencing data from normal brain^16,39^, breast^17^, and colon samples^40^ (Supplementary Table 7). Intriguingly, breast tissue exhibited a significantly higher proportion of such mutations compared to the other two tissue types (p=0.0012, two-tailed t-test, Extended Data Fig. 11a-b). Notably, the vast majority of these mutations (44/45) were found within B cell receptor (BCR) genes. However, only a subset of these BCR mutations (16/44, ∼36%) were found in hypervariable regions typically associated with somatic hypermutation. This distribution implies that although clonally expanded B cell populations are present in normal breast tissues, their mutational signatures may involve mechanisms extending beyond classic antigen-driven affinity maturation. Interestingly, the BCR mutations mapping to variable regions exhibited significant spatial clustering across normal breast samples (Extended Data Fig. 11c-d). This pattern suggests focal, antigen-driven selection pressures, potentially reflecting compartmentalized immune activity that could contribute to tissue-specific immunosurveillance or early alterations in the tissue microenvironment. Functional investigation of these expanded B cell clones may clarify their role in breast homeostasis and pathogenesis, offering insights for risk assessment or immunopreventive strategies.

## Discussion

The development of SpaceTracer marks a significant advancement in the field of spatial lineage tracing. By integrating lineage information, spatial organization, and gene expression profiles, our framework enables a unified and comprehensive view of complex biological processes *in vivo*. This integrated approach is uniquely positioned to reveal the spatiotemporal dynamics underlying development, homeostasis, and disease progression.

The key strength of SpaceTracer lies in its ability to reconstruct lineage relationships directly from naturally occurring somatic SNVs. This feature is particularly critical for human studies, where the genetic manipulations routinely used in model systems are not ethically permissible. Furthermore, unlike profile-based approaches, somatic SNV occurrence is a binary (“zero or- one”) event. This “digital” nature enables the spatial tracing of minority cell populations even at multicellular resolution—a task fundamentally infeasible with gene-expression-based methods, which lack intrinsic accuracy for precise cellular parsing and tracking. By integrating information from cells within the same lineage, a coupled analysis of somatic mutations and expression profiles simultaneously defines cellular identity and lineage history. Even when cells from a given lineage migrate into spatial domains dominated by other cell types, they remain traceable via their unique mutations, regardless of how low their abundance becomes in the new niche.

SpaceTracer is particularly advantageous for decoding tumor progression and tumor-immune interactions, leveraging the abundant mutations and clonal expansions in tumors and immune cells. When applied to cSCC, it uncovered pivotal dynamics, such as the widespread migration of non-tumor-forming epithelial cells across two independent donors. Expression analysis revealed these cells probably had undergone dedifferentiation—an ancestral reversion that likely fuels early tumor initiation and invasion. We also traced mutant B cells as they migrated from TLS to infiltrate the tumor boundary. Given the abundance of hypermutations within TLS regions, SpaceTracer also offers a powerful, perturbation-free method to characterize these structures and study the behavior of mutated B cells—insights that hold promise for advancing tumor immunotherapy.

Taken together, SpaceTracer establishes a generalizable framework for dissecting tissue dynamics at unprecedented resolution. As spatial multi-omics datasets grow in scale and complexity, our tool is poised to become increasingly vital, with transformative potential across biomedicine: from deciphering tumor heterogeneity and therapeutic resistance in oncology to mapping cellular fate decisions and interactions in development and aging. SpaceTracer is designed for the broad research community for accessibility, integrating accurate mutation detection, lineage reconstruction, and the generation of publication-ready figures in a single, user-friendly environment. While currently optimized for nuclear SNVs, future iterations will incorporate mitochondrial mutations and additional omics layers (e.g., epigenetics), further advancing high-resolution, multimodal lineage tracing in vivo.

## Acknowledgements

We are deeply grateful to the generous tissue contributions. We acknowledge Drs. Hongtao Yu, Jiyuan Yang, Ting Zhou, Jinran Lin, Bing Zhang, Jianyang Zeng, Weike Pei, Yanxiao Zhang, Dangsheng Li, Shang Cai, Qi Xie, Heping Xu, Min Jiang, Li Li, Shouwen Wang and Jian Yang for their invaluable insights and enriching discussions. We would want to acknowledge Dr. Andrew L. Ji for sharing the cSCC data. We thank the Flow Cytometry Core Facility and the Genomics Core Facility at Westlake University for their experimental assistance, and the High-Performance Computing Center for technical support. This work was supported by the Westlake Laboratory of Life Sciences and Biomedicine (Hangzhou 310024, Zhejiang, China), under the grant “Key R&D Program of Zhejiang” (2024SSYS0034) as well as the National Natural Science Foundation of China (32270682) to Y.Dou. We also acknowledge financial support from the Westlake Education Foundation, as well as from the Research Program No. WU2023C020 of Research Center for Industries of the Future, Westlake University. We would like to thank the submitters of the following datasets: GEO accession GSE158328, GEO accession GSE144240, GEO accession GSE225475, GEO accession GSE220978, GEO accession GSE237183, GEO accession GSE195665, GEO accession GSE171351, GEO accession GSE176078, GEO accession GSE274641 and GEO accession GSE161363. These public datasets were obtained from GEO at https://www.ncbi.nlm.nih.gov/geo/. And the accession number EGAS00001006124 from the European Genome-phenome Archive (EGA) at https://ega-archive.org/; the accession ids (jhpce#spatialDLPFC and jhpce#HumanPilot10x) from Globus at https://www.globus.org/.

## Author contribution

Y.Dou conceived and supervised the study, Y.Dou and Y.Z (Weslake University) acquired funding. H.Y. assisted in supervision. Z.Y., M.Y., Q.Y., Y.Du, and J.Lu contributed equally. Z.Y. and M.Y. developed software and performed data analysis. Q.Y. conducted phylogeny reconstruction, Y.Du. contributed to data analysis, and J.Lu carried out software benchmarking. J.Lin, Y.Z (Huashan Hospital Fudan University), J.Luan, Q.W., Q.Z., J.Lu, X.M., Y.L. were involved in sample collection and wet-lab experiments. X.W., Z.Q., S.H., J.Lu, Q.Z., H.L., W.F., Yuhao Xia and Yonghe Xia either performed or assisted with experiments and data analysis. Y.Dou, Z.Y., M.Y., Y.Du and J.Lu wrote the manuscript. All authors carefully reviewed and approved the final manuscript.

## Inclusion & Ethics statement

The authors affirm that the research was conducted in accordance with ethical guidelines and principles. All data used in this study are publicly available, and no personally identifiable information was used.

## Code availability

SpaceTracer is implemented in Python and R and is licensed under the MIT License. The SpaceTracer source code is being prepared for public release. It will be available on GitHub (https://github.com/douymLab/SpaceTracer) by March 1, 2026, or upon the manuscript’s formal acceptance, whichever comes first.

## Data availability

WGS data and spatial transcriptomics data generated in this study will be publicly available in the Sequence Read Archive (SRA) (PRJNA1396783, Link: https://dataview.ncbi.nlm.nih.gov/object/PRJNA1396783?reviewer=m2j7hed6okvjj82stkthdue2te). The published spatial transcriptomics data and single-cell RNAseq data analyzed in this study could be obtained from GEO at https://www.ncbi.nlm.nih.gov/geo/ with GEO accessions GSE158328, GSE144240, GSE225475, GSE220978, GSE237183, GSE195665, GSE171351, GSE176078, GSE274641 and GSE161363. And the prostate cancer data were obtained by the accession number EGAS00001006124 from EGA at https://ega-archive.org/. The normal brain data were obtained from Globus (accession ids (jhpce#spatialDLPFC and jhpce#HumanPilot10x)) at https://www.globus.org/. See detailed data resources in Supplementary Table 2.

## Methods

### Sequencing data processing

#### Pre-processing of whole-genome sequencing data

The whole genome sequencing data were aligned to the GRCh38 reference using BWA-MEM (v0.7.17), and duplicates were detected and removed using GATK MarkDuplicates (v4.2.6.1).

#### Pre-processing of single cell sequencing data

The scRNA and snRNA data collected from public dataset were downloaded in fastq format and aligned to GRCh38 build of the human reference genome by CellRanger^1^ (v7.1.0). The procedures for dimensionality reduction, and clustering were executed through Seurat (v5.1.0). For clustering, we applied Seurat’s LogNormalize method with 2,000 variable features by default. Then, the data was scaled after regressing out mitochondrial gene percentages. And the nearest neighbor search was conducted using the RANN method, and principal component analysis (PCA) was performed, retaining the first 30 principal components.

#### Pre-processing of spatial-transcriptomic sequencing data

The spatial transcriptomic Visium data of human were aligned to the GRCh38 reference, the same reference as the single-cell data, using SpaceRanger (v2.0.1). All output BAM files were processed uniformly. First, we required that each read have no more than 5 mismatches and that the mapping quality (MapQ) of each read be greater than or equal to 255. The filtered reads were then used for subsequent mutation detection.

### SpaceTracer workflow

#### 1. Prescan and quality filtering of candidate mutations

VAFs were called from the filtered BAM files using samtools mpileup (v1.21) (samtools mpileup --excl-flags 0 -s -B -Q 0 -q 0 -d 200000 -f reference.fa sample.bam). Candidate mutations with low total read depth (<30×), low alternative read depth (<5×), high VAF (>0.3) and low VAF (<0.1%) were excluded. Population allele frequencies for the corresponding alleles were then retrieved from the gnomAD database. To identify and exclude likely sequencing artifacts, we selected candidate mutations with a VAF < 5% in our sample and a population allele frequency of 0% in gnomAD. The median noise level (0.3%) was determined as the empirical background error rate, and candidate mutations with their VAFs not significantly higher than 0.3% were excluded (one-tailed binomial test, p>0.05).

#### 2. Reads aggregation at the UMI level

To minimize artifacts from over-sequenced UMIs, we aggregate reads at the UMI level before genotyping:

◯ The posterior probability that allele j generates reads with UMI identifier “*umi* − *k*” can be calculated by Bayes rule as:

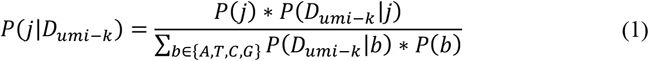 Where:
  ▪ *D*_*umi−k*_ = reads with UMI identifier “*umi* − *k*”,
  ▪ *b* = all possible alleles, including A, T, C, G,
  ▪ *P*(*j*) = prior probability of j allele, which equals to ¼, same for *P*(*b*).
◯ Accounting for sequencing errors or PCR errors, the likelihood of observing reads with UMI ID “umi-k” given j allele is approximated by: 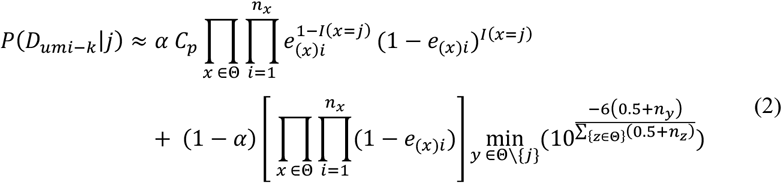 Where:
  ▪ α indicates the relative weight between observing a true allele (or an allele resulting from sequencing errors, as captured in the first part of the formula) and observing an allele due to PCR errors (as described in the second part of the formula). The default value of α is set to 0.5.
  ▪ *e*_(*x*)*i*_ indicates sequencing error, where 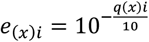.
  ▪ θ is the set of {A,T,C,G}.
  ▪ *n*_*y*_ indicates the number of y alleles in the sequencing reads, so for *n*_*z*_.
  ▪ 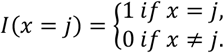
◯ The “consensus allele” *j* for reads with barcode *umi* − *k* was defined as the allele with the highest posterior probability. Its combined error rate is given by: *E*(*j*) = 1 − *P*(*j*|*D*_*umi−k*_)

#### 3. Hierarchical Bayes genotyping

For genotyping, reads with >5 mismatches or mapping quality <255 were filtered. The hierarchical genotyping likelihoods are then calculated based on UMI-level allele counts as follows. We use Bayes rule to evaluate the individual-level and spot-level genotype with four genotypes: heterozygous (het), reference-homozygous (refhom), alternative-homozygous (althom) and somatic mutation (mosaic). With *G* denotes the genotype and *D* denotes the data, the Bayes rule could be represented as below.

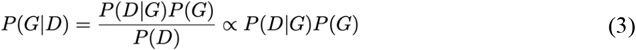

The prior for each genotype is estimated by the population allele frequencies, which could be achieved from dbSNP (the database of Single Nucleotide Polymorphisms)^2^ and The Genome Aggregation Database (gnomAD)^3^. The formulas are shown below:

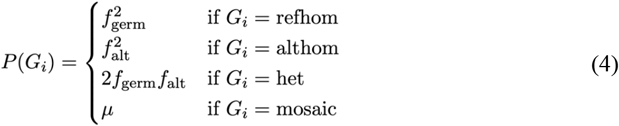

Where *f*_germ_ and *f*_alt_ are the population allele frequency of the germline and alternative allele at the corresponding site respectively. µ approximates the population somatic mutation rate, we would use µ = 10^−7^/bp as a default in our algorithm.

The theoretical mutant allele fractions (MAFs) for each genotype are 0, 0.5, and 1 for refhom, het and althom respectively. Assuming the MAF for mosaic genotype is uniformly distributed between 0 and 1, we used Bernoulli sampling to approximate the PCR amplification error, sequencing errors and the bias between the reference and alternative allele read depth^4^. The likelihood functions for the four genotypes hence could be computed by the following equations:

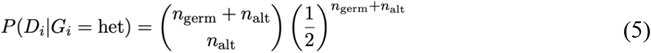

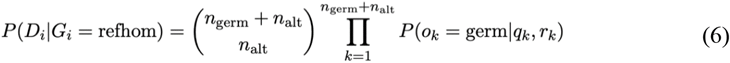

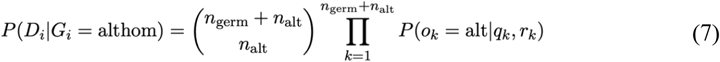

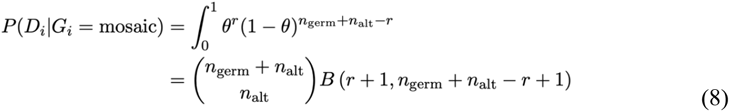

Where:

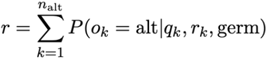

- *n*_germ_ is the number of consensus reads (with the same UMI) of the germline allele;
- *n*_alt_ is the number of consensus reads (with the same UMI) of the alternative allele;
- *r*_*k*_ is the consensus read *k*;
- *o*_*k*_ is the observed allele on consensus read *k* at the mutant position;
- *q*_*k*_ is the quality on the consensus read *k* at the mutant position;
- θ is the real mutant allele fraction;
- *G*_*i*_ denotes the individual genotype;
- *D*_*i*_ denotes the observed individual data;
- *B* denotes the beta function.

We hypothesize that spots within solid tissues that are in close proximity, exhibit similar expression patterns, or correspond to the same cell type are more likely to share the same genotype. Based on this assumption, we divided the spots into clusters considering both gene expression and location information and assumed that spots within the same cluster are more likely to exhibit similar mutant allele frequencies (VAF). SpaGCN (v1.2.7)^5^ and GraphST (v1.1.1)^6^ are the default clustering algorithms used in our approach, as they account for both gene expression levels and spatial locations when grouping spots. After the initial clustering, density-based spatial clustering of applications with noise (DBSCAN) implemented in the scikit-learn (v1.2.2) was used to separate disconnected clusters into distinct domains^7^. In cases where small domains contain only 1-2 spots, we merge them with adjacent domains that are part of the same cluster.

The prior probabilities were then calculated by considering the mutant allele frequency of the cluster, as shown below:

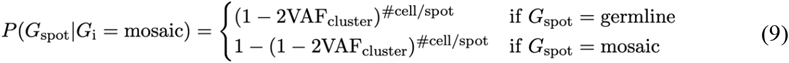

Here, VAF_cluster_ = AF_mutant allele,cluster_, where the allele frequency of the mutant allele is determined for the cluster to which the spot belongs. Additionally, “#cell/spot” denotes the number of cells per spot. As in most cases, spatial transcriptomics data contains spots with more than one cell. The likelihood functions for the spots are identical to those of the individual likelihood functions. Set the default threshold to 0.5, allowing us to identify a spot as mutated if its mosaic genotype posterior exceeds 0.5 and it contains at least one mutant allele.

#### 4. Comprehensive feature extraction

- SpaceTracer incorporates 98 features (Supplementary Table 1) spanning:
  - **Read-level statistics**: mapping quality difference, read alignment positional bias, base quality profiles, base position distribution within reads, per-read mismatch counts, indel-containing read fraction, soft-clipped read proportion, homopolymer-related filtration, etc.
  - **RNA-specific metrics**: strand bias assessment, RNA editing potential (filtered against RADAR, REDIportal, and DARNED), UMI-level allele concordance, gene-expression-level correlated artifacts, allele-specific expression (ASE) analysis, etc.
  - Spatial-**distribution related features**: Kolmogorov–Smirnov (KS) tests^8^ comparing the spatial distributions of mutant and non-mutant spots and Moran’s I index^9^ quantifying spatial autocorrelation of mutant spots.
- Standardization for depth-invariant features:
  - To ensure robust feature comparisons between reads carrying reference and alternative alleles, we employed standardized effect sizes rather than raw test statistics. For example, to address depth heterogeneity, we standardize Wilcoxon rank-sum test statistics (U) by calculating the Rank-Biserial Correlation (RBC) for feature comparisons (e.g., alt vs. ref reads): 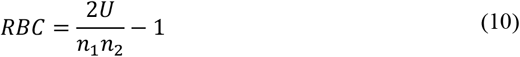 Where:
    ▪ *U* stands for the test statistic value in the Wilcoxon rank sum test,
    ▪ *n*_1_ and *n*_2_ are the sample sizes for the two groups (e.g. the number of reference and alternative alleles).
  - Moreover, before feature extraction, we apply a depth-capping procedure to each candidate locus. If the sequencing depth is greater than 2000×, reads are randomly down sampled to exactly 2000×; otherwise, all available reads are used. This step is essential in single-cell RNA-seq analysis to prevent highly expressed loci from dominating the features and to mitigate bias arising from the vast differences in sequencing coverage.

#### 5. Somatic mutation prediction using a machine learning model based on a rigorously curated training set

- Rigorously curated training set with orthogonal validation:
  - We constructed a high-quality benchmark dataset for model training and testing, comprising over 18 slides from seven independent individuals. Each public spatial dataset is paired with at least two orthogonal data sources for cross-validating candidate mutations—such as matched scRNA-seq data or additional spatial transcriptomics data from adjacent tissue slices. A “leave-one-donor-out” cross validation strategy was used. To ensure robust validation, we generated a benchmark data by performing deep whole-genome sequencing (WGS; ∼200× coverage) on adjacent tissue slices of a BCC sample. This comprehensive, real-world dataset, reinforced by multi-modal cross-validation, allows our model to learn biologically accurate parameters.
  - Two models were trained: a spatial-preserved model and a spatial-feature-free model. This design allows for somatic mutation prediction across datasets where clonal populations either maintain or lack clear spatial organization, as in cases of extensive cell migration. The spatial-feature-free model was trained and evaluated using all available samples. During its training, the following five spatial-distribution features were explicitly removed: “pass_spatial_test”, “mut_vs_nonmut_spots_KS_p”, “mut_vs_nonmut_spots_KS_s”, “mut_vs_nonmut_spots_MI_p” and “mut_vs_nonmut_spots_MI_s”. In contrast, the spatial-preserving model was trained exclusively on a curated subset of datasets exhibiting definitive spatial organization: two skin cancer datasets and a breast cancer dataset (see Supplementary Table 2).
- Training set construction facilitated by haplotype phasing:
  - The low in vivo somatic mutation rate makes it highly impossible to for a mutation to occur at the same genomic position on both paternal and maternal haplotypes. We utilized this property by using *in silico* haplotype phasing^10^ to guide the training set construction. True somatic mutations are expected to be present in only one haplotype within a subpopulation of cells. In contrast, mutations appearing on both haplotypes across cells are likely artifacts caused by mis-mapping. In our training set, ∼96% of artifact mutations were identified based on this phasing information. The left artifact mutations were identified following an extensive manual review process, as described in the “Cross-validation of mutations using sequencing data from adjacent slices” section.
  - To obtain an unbiased performance estimate, we adopted a leave-one-donor-out cross-validation strategy. Specifically, for each donor designated as the test set, all samples from that donor were excluded from the training data, and a model was built using samples from all other donors only. This ensures that the model evaluation reflects generalizability across independent individuals.

#### 6. Further filtration of likely lysis errors and other chemical modifications in RNA utilizing artifact mutation signature analysis

While library preparation artifacts (e.g., chemical modifications) and RNA-editing events may mimic true somatic mutations at the read-level, their mutational patterns often exhibit distinct signatures. To systematically identify and exclude these artifacts, we did:

#### Artifact signature deconvolution

- Extracted 192-category RNA mutation profiles (accounting for trinucleotide context and strand-specificity) from ultra-low-frequency variants (defined as those supported by only one UMI with allele frequency <5%).
- Identified three dominant artifact signatures by applying non-negative matrix factorization (NMF, v0.28)^11^ to low VAF variants from 101 slices of 50 individuals (Supplementary Table 2: data used for signature). The optimal number of signatures was determined with Variational Bayesian Factorization using ccfindR (v1.18.0)^12^, balancing model fit and overfitting avoidance.
- The major artifact patterns are characterized and detailed the “Artifact signature deconstruction” section (Extended Data Fig. 2):
  - C>T transitions: Consistent with heat-induced cytosine deamination during lysis.
  - G>T transversions: Indicative of oxidative DNA damage during in vitro processing.
  - A>G mutations: Likely reflecting RNA-editing processes.

Quantitative Filtering:

- For each candidate mutation we derived a heuristic false discovery rate (hFDR) score incorporating and excluded candidates with hFDR ≥ 0.8 to minimize technical false positives (Extended Data Fig. 2):

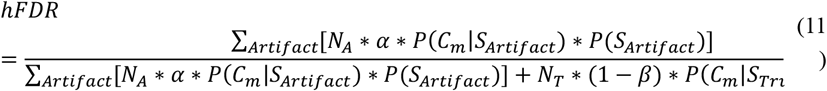

Where:

- α and β denote type I and type II errors discovering no less than *k* mutant reads for mutation *m*;
- *S*_*Artifact*_ = (*S*_*A1*_, *S*_*A2*_, …, *S*_*A1*92_), representing the 192-channel mutational signature of a specific artifact;
- *S*_True_ = (*S*_T1_, *S*_T2_, …, *S*_T192_), representing the 192-channel mutational signature of true somatic mutations, *S*_T1_ = *S*_T2_ = … = *S*_T192_ = 1/192;
- *P*(*C*_*m*_|*S*_*Artifact*_) denotes the likelihood of observing a mutational channel *C*_*m*_ given the artifact signature *S*_*Artifact*_, *P*(*C*_*m*_|*S*_*True*_) is defined similarly for the true signature;
- *P*(*S*_*Artifact*_) is the probability of an artifact signature given the data from a specific sample, estimated using Non-Negative Matrix Factorization (NMF); *P*(*S*_*True*_) defined in the same way for true somatic mutations; ∑_*Artifact*_ *P*(*S*_*Artifact*_) = *P*(*S*_*True*_);
- *N*_*A*_ and *N*_*T*_ represent empirical number of artifact mutations and true somatic mutations,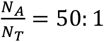.

The distribution of background noise, represented by the alternative allele fraction, is modeled by a beta distribution with shape parameters a and b:

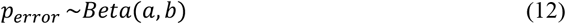

We assume a uniform prior distribution for the empirical alternative allele fraction (AAF) at true somatic SNVs over the interval [0, 1]:

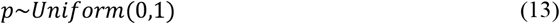

Then the type I and type II errors discovering no less than *k* mutant reads for mutation *m* could be calculated as follows:

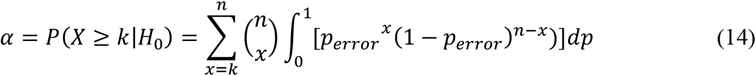

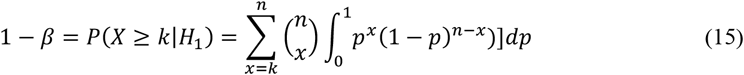

Where:

- *n* denotes the total read depth,
- *x* denotes the alternative allele count,
- *k* denotes the observed alternative reads.

#### 7. Further eliminate (recurrent) artifacts using a series of hard filters

- Population variant database filtration: variants present in the gnomAD database with >0.01% population allele frequency were excluded.
- Filter variants in genes with allele-specific expression (ASE): A gene was classified as exhibiting ASE if it contained at least one likely heterozygous germline variant (with the highest genotyping posterior supporting a germline call and a gnomAD population AF >0.01%) with significant allelic imbalance (two tailed binomial test, p≤0.05). Within these genes, candidate somatic variants whose variant allele frequency (VAF) was not significantly different from a nearby germline SNP’s allele fraction (two-tailed binomial test, p>0.05) were excluded as potential false positives.
- RNA Editing Exclusion:
  - Remove all RNA-editing mutations annotated in three major RNA-editing databases: RADAR^13^, REDIportal^14^, and DARNED^15^.
  - All A>G mutations were removed, as these are likely RNA-editing events not excluded by standard RNA-editing databases.
- Remove mutations within imprinted genes: mutations within 227 classical imprinted genes were (http://www.geneimprint.com/) and excluded.
- Mutations within or nearby homopolymers (6bp or longer) were excluded.
- Remove variants present in panel of normals: SpaceTracer uses a panel of normal generated with 33 non-neoplastic samples to discount recurrent artifacts and germline variants.

### Mutation signature analysis across tissues

Tissue-specific 96-category mutational profiles were extracted and plotted using SigProfilerAssignment (v0.2.6)^16^. Extracted signatures were compared to known COSMIC signatures with deconstructSigs (v1.9.0)^17^. To avoid overfitting, comparisons were restricted to signatures whose characteristic mutational peaks are observed in empirical data: SBS1, SBS5, SBS7a, SBS7b, SBS7c, SBS7d, SBS38 for skin; SBS1, SBS2, SBS5, and SBS18 for brain and SBS1, SBS5, SBS40 for breast.

#### Mutation rate estimation

- In single cell sequencing data, the mutation rate per cell µ_*c*_ can be estimated by dividing the number of detected mutations in a cell by the size of its callable genomic region:

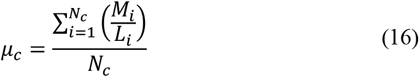 Where:
  - *M*_*i*_: the total number of mutations detected in the ith cell.
  - *L*_*i*_: the total length (bp) of the callable genomic region for the ith cell.
  - *N*_*c*_: the total number of cells analyzed.
- In spatial transcriptomics data, the single-cell level barcoding is unavailable. The mutation rate per cell could be approximated with a bounded interval: 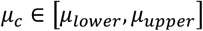

Where the lower and upper bounds are defined as:

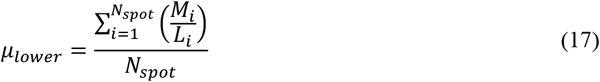

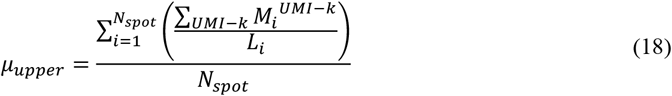

Where

◯ *UMI* − *k*: the kth UMI in spot *i*.
◯ *M*_*i*_: the total number of mutations detected in spot *i*.
◯ *L*_*i*_: the size (bp) of the callable genomic region for spot *i* (with non-zero coverage in at least 30 spots and non-zero coverage in spot *i*).
◯ *N*_*Spot*_: the total number of spots analyzed.

The lower bound (*µ*_*lower*_) represents the most conservative estimate, valid if all mutations with distinct UMIs within a spot originate from a single cell. The upper bound (*µ*_*upper*_) represents the maximum estimate, valid if each mutation with a unique UMI originates from a distinct cell.

In practice, due to the high data sparsity of spatial transcriptomics data (∼60-80% of mutations are supported by only a single UMI), the values of *µ*_*lower*_ and *µ*_*upper*_ are highly constrained (typically differing by only ∼10–20%). Consequently, we employ the more conservative and stable lower-bound estimate (*µ*_*lower*_) for all subsequent analyses.

Moreover, to estimate the mutation rate in normal-skin regions using cSCC samples, we distinguished between normal and tumor tissue based on the TSK score and leading-edge assessment (see “Designation of clonal regions for mutation rate estimation” for more details). For each slice, mutation burden was computed separately for the normal and tumor regions, which were then annotated as normal skin and tumor skin, respectively, for subsequent analysis.

### Published mutate rates derived from DNA-seq data

The upper and lower bounds of mutation rate per cell for each tissue were derived from recent literature as follows:

- Tumor Skin: 0.1–380 mutations per Mb, based on deep targeted DNA sequencing of cSCC and BCC samples^18^.
- Normal Skin: 0.4–12 mutations per Mb, derived from deep targeted DNA sequencing of sun-exposed facial skin in Singaporean and UK populations^19^. The donor age range (68-90 years) is consistent with that of our benchmark cohort (74–96 years).
- Normal Brain: 0.167–1.3 mutations per Mb (estimated from single-cell whole-genome sequencing data of age-matched individuals, 20–60 years)^20^.
- Normal breast: 0.1–0.4 mutations per Mb (estimated from WGS data of single-cell oganoids from age-matched donors <60 years)^21^.

### Benchmark of SpaceTracer

#### Mutation detection using different software tools

We compared the performance of SpaceTracer against three algorithms: SComatic^22^, Monopogen^23^ and SpatialSNV^24^. Scomatic (vfrom 2024-01-10) was run with n_trim=5, max_nM=5, max_NH=1 in the first step, min_bq=30 in the second step, all other parameters are set to the default values. For Monopogen (v1.5.0), we run the data preprocesses, germline SNV calling and putative somatic SNV calling with default values for all parameters, then filter the putative SNVs with the suggested hard filters: SVM_pos_score>0.5, LDrefine_merged_score>0.25, 0.1<BAF_alt<0.5, Dep_ref>2 and Dep_alt>2. For SpatialSNV (v1.1.3), first, input BAM files were split by chromosome using the spatialsnvtools SplitChromBAM function to enable parallelization. Second, mutation calling data was preprocessed with spatialsnvtools PrepareBAMforCalling. The SNV callings were then produced based on the preprocessed files using the spatialsnvtools SNVCalling function. MNVs were excluded. All steps were executed with default parameters. In the tested samples, SpatialSNV typically identifies tens of thousands of mutations per sample, the vast majority of which are at very low allele frequencies. To generate a high-confidence callset, we applied additional filters requiring a minimum of 30 spots with non-zero coverage, a minimum alternative allele count of ≥3 supporting UMIs, and excluded likely germline variants with VAFs ≥ 30%.

#### Sample collection and sequencing

Samples were collected at Huashan Hospital, Fudan University. After surgical resection from patients with basal cell carcinoma, the specimens were immediately immersed in a Tissue Storage Solution (Miltenyi, 130-100-008), placed on ice, and transported to Accuramed Biotechnology Co., Limited.

Fresh tissue specimens were embedded in Optimal Cutting Temperature (OCT) compound (SAKURA) and stored at -80°C until use. Immediately after collecting ten frozen sections with a thickness of 10 μm, RNA integrity was assessed using the RNeasy Mini Kit (Qiagen). Samples with an RNA Integrity Number (RIN) ≥7 were considered qualified for subsequent experiments.

Tissue permeabilization was optimized for the 10 μm-thick sections using the Visium Spatial Tissue Optimization Kit (10x Genomics). In this study, the optimal permeabilization time varied from 3 to 30 minutes depending on the specific sample. Qualified sections were mounted onto Visium Spatial Gene Expression Slides (10x Genomics), fixed with methanol (Millipore Sigma), and stained with Hematoxylin and Eosin (H&E). The stained slides were then imaged using a Leica sc2 slide scanner (Leica Biosystems). The tissue sections were permeabilized to release polyadenylated mRNA, allowing capture by the pre-coated poly(dT) primers on the slide. These primers contained sequences for the Illumina TruSeq Read 1, a Spatial Barcode, and a Unique Molecular Identifier (UMI).

The Visium Spatial Gene Expression Reagent Kit (10x Genomics) was used for reverse transcription to generate spatially barcoded, full-length cDNA, followed by second-strand synthesis. The cDNA was then denatured and transferred from the slide to tubes for amplification and library construction. The quality and quantity of the synthesized cDNA were assessed by running the sample on an Agilent Bioanalyzer High Sensitivity chip (Agilent).

Visium spatial gene expression libraries, comprising P5, P7, i7, and i5 sample indexes along with the TruSeq Read 2 primer sequence, were constructed through enzymatic fragmentation, end repair, A-tailing, adaptor ligation, and sample index PCR. Unique i7 and i5 sample indexes were added using the Dual Index Kit TT Set A (10x Genomics). Finally, the constructed libraries were sequenced on an Illumina NovaSeq 6000 system.

This experiment was performed following the standard operating procedure of the 10x Genomics Visium Spatial Gene Expression technology platform. Unless otherwise specified, all core reagents and consumables were sourced from this platform, primarily including: the Visium Spatial Gene Expression Slide & Reagent Kit (4 reactions, PN-1000187), Visium Accessory Kit (PN-1000194), Visium Slide Cassette & Gasket (PN-2000281), and the 10x Magnetic Separator (PN-230003). All steps were strictly performed according to the manufacturer’s instructions (User Guide CG000239, Rev H).

#### Cross validation of mutations using deep WGS sequencing of DNA

In our benchmark dataset (BCC-1), two adjacent tissue sections were selected from consecutively sectioned, OTC-embedded samples, with one section used for spatial transcriptomic and the other used for whole genome amplification (WGA). For the WGA section, tissues were mounted on MMI PPS Membrane Slides (MMI, 50112) and subjected to hematoxylin and eosin (H&E) staining using slide stainer (LEICA, ST5020). Normal and tumor regions were identified by clinical experts based on the staining results. These regions were subsequently isolated by Laser Microdissection System (MMI, CellCut Plus), and the normal and tumor tissues were collected separately into MMI Isolation Caps transparent (MMI, 50208) and subjected to WGA using the ResolveDNA kit (Bioskryb). The amplified DNA was then subjected to PCR-free, deep whole-genome sequencing (WGS) to an average coverage of ∼200×.

Somatic mutation calling was performed separately for normal and tumor samples using Mutect2^25^ (gatk v4.2.2.0) with a panel-of-normals (PON) mode. The resulting variant calls were further filtered using MosaicForecast^26^ with default parameters. To ensure that the identified whole-genome somatic mutations could also be detected in the transcriptomic data, we required each candidate site to meet the following data-quality thresholds: at least 30 UMI-covered spots, a minimum of 3 alternative-supporting UMIs, and an alternative allele fraction (VAF) ≤ 0.3 These parameters were also used as filtering criteria in SpaceTracer.We used samtools (v1.21) mpileup with stringent quality filters (-s -B -Q 20 -q 30 --excl-flags SECONDARY,QCFAIL -d 200000) to extract high-quality allele depths. To validate mutations, we employed a two-step approach: 1. Statistical validation of mutant alleles: We performed a one-tailed Fisher’s exact test to compare the read count supporting the top mutant allele with that of the second-highest allele. Mutations were considered validated at a significance threshold of *p* ≤ 0.05. 2. Exclusion of likely germline variants: Putative somatic mutations were defined as those with a variant allele frequency (VAF) significantly less than 50%, as assessed by a one-tailed binomial test (*p* ≤ 0.05).

#### Cross validation of mutations using sequencing data from adjacent slices

Candidate mutations identified by different software tools in public benchmarks—including tumor skin (cSCC P4, P6), normal breast (P10), and normal brain cortex (Br5292, Br5595, Br8100) samples—were cross-validated using a multi-tiered verification approach. This involved: (1) manual inspection of read alignments by independent researchers; (2) statistical testing for cell-type distribution biases; and (3) confirmation using matched data from adjacent slices.

First, all mutations called by the software tools underwent extensive manual review by three independent researchers. We generated read-alignment plots at both the spot and pseudo-bulk levels using IGV^27^ (v2.18.2). At the spot level, spots were separated into two groups based on the presence of alternative allele reads. A candidate mutation was classified as an artifact (False) if IGV inspection revealed typical artifactual features. These included: a pronounced end-enrichment pattern of ALT reads, an excessive number of mismatches within ALT-supporting reads, or location within a homopolymer of ≥6 bp.

Second, for the public benchmarks where matched scRNA-seq data from adjacent tissue was available (cSCC P4, cSCC P6, and normal breast P10), candidate mutations that passed manual inspection were classified as “validated somatic” based on the following criteria: 1) the mutation was detected in at least one single cell in the scRNA-seq data, and 2) its cell-type distribution was statistically assessed. Specifically, for mutations with a variant allele frequency (VAF) >10% in the scRNA-seq data, we performed a Fisher’s exact test. Mutations that did not show a significant difference in the proportion of mutant cells across distinct cell types (p > 0.05) were considered putative germline variants and were excluded. For the public benchmarks where spatial transcriptomics data from adjacent tissue slices were available (Br5292, Br5595, Br8100), candidate mutations that passed manual inspection were classified as “validated somatic” if they were detected in at least one spot in those adjacent slices. Otherwise, the mutation will be marked as “undetermined”.

#### Benchmark of phylogeny reconstruction

To establish a phylogeny benchmark, we leveraged two public lung adenocarcinoma (LUAD) scRNA-seq datasets harboring knock-in CRISPR barcode mutations^28^. These single-cell datasets were spatially projected using CytoSpace^29^ (v1.1.0) to generate two corresponding simulated spatial transcriptomics datasets, each possessing a known ground-truth barcode phylogeny. Specifically, cells from the scRNA-seq benchmarks were mapped onto the spatially resolved spots of a publicly available human LUAD spatial transcriptomics dataset (10x Genomics Visium HD, https://www.10xgenomics.com/datasets/visium-hd-cytassist-gene-expression-human-lung-cancer-fixed-frozen). To ensure a strict one-to-one correspondence between cells and spatial spots, we configured CytoSpace with the restriction that each cell from the scRNA-seq dataset could be mapped to only one spatial spot, and that each spatial spot could harbor at most one scRNA-seq cell. This mapping procedure thereby conferred spatial context to the single-cell data, creating a realistic simulated spatial benchmark with defined phylogenetic ground truth.

Somatic mutations from the simulated benchmarks were initially identified from the scRNA-seq data using MosaiSC^18^. Spatial genotyping was subsequently performed using SpaceTracer. Phylogenetic trees were then reconstructed with PhyloSOLID (v1.0.0)^30^, using all default parameter settings.

### CNV calling

CNVs were inferred from cSCC-1 with two state-of-the-art tools: a reference-based method inferCNV (v1.20.0)^31^ and a reference-free, allele-aware method CalicoST (vfrom2024-12-10)^32^.

To construct the reference for inferCNV, we first identified reference spots based on cellular deconvolution results. These were defined as spots predominantly composed of non-keratinocyte and non-epithelial cell types, as detailed in the “Single-cell transcriptomics annotation and spatial transcriptomics deconvolution” section. Using these reference spots as a baseline, we ran inferCNV with default parameters on the gene expression count matrix. The algorithm generated a matrix of inferred relative copy number segment ratios. These continuous predictions were then discretized into six states, which we consolidated into three broad categories: copy number loss (states 1 and 2), neutral (state 3), and copy number gain (states 4–6). For subsequent integrative analysis, all non-neutral states (1, 2, 4, 5, 6) were collectively termed “copy changed.”

We performed copy number analysis using CalicoST following its standard protocol. First, we executed the preprocessing module to generate allele-specific count matrices after genotyping and reference-based phasing. Second, we applied the estimate_tumor_proportion module to infer tumor purity per spot. Third, using these purity estimates as input, we ran the main CalicoST module to infer subclonal copy number aberrations, configured to identify up to three major subclones (n_clones = 3), with default parameters. All analyses utilized the GRCh38 reference genome.

### Functional and genomic region annotations

The functional and genomic region annotation of identified mutations was performed using ANNOVAR^33^. Gene-based annotation was conducted with GENCODE (v46) gene definitions, obtained from the UCSC Genome Browser. For non-synonymous variants, functional predictions were derived from the dbNSFP database within ANNOVAR, which includes Polyphen2^34^ scores based on the HumanDiv.

### Pathway enrichment analysis

We conducted functional enrichment analysis with KOBAS (v 3.0.3)^35^. The analysis covered: (1) biological pathways from KEGG and Reactome databases, (2) disease-associated pathways from KEGG Disease, and (3) functional annotations from Gene Ontology (GO).

### Cell type deconvolution and annotation

To construct a comprehensive cell-type reference atlas for deconvolution, we collected published scRNA-seq datasets from matched tissues (Supplementary Table 6). Using this atlas as a reference, we then annotated cell types within our spatial transcriptomics data with Spotify (v0.1.1)^36^. For cSCC samples lacking pre-defined annotations in the original scRNA-seq datasets, we performed *de novo* cell-type annotation by projecting them onto the Blueprint and ENCODE reference database using SingleR (v2.0.0)^37^.

The deconvolution process with spotify was conducted following the recommended tutorials. First, the spotiphy.deconvolution.estimation_proportion function (n_epochs=8000) was used to estimate the relative proportion of each reference cell type within every spatial spot. Next, we estimated the absolute cell number per spot using an in-house pipeline. This estimation is based on the assumption that cell count is positively correlated with a spot’s total UMI count. We applied a linear regression model to this relationship. Given the spot diameter of 55 *µ*m, the model was constrained such that the maximum predicted cell number per spot could not exceed 25. We then supplied the estimated absolute cell numbers per spot and the relative cell-type proportions to the spotipy.deconvolution.simulation module. Using these inputs under preset parameters (batch_effect_sigma=0.1, zero_proportion=0.3, additive_noise_sigma=0.05), the module generated simulated counts, from which the approximate cell count per cell type per spot was calculated.

### Tumor specific keratinocytes (TSK) and tumor fitness signature score calculation based on gene expression

TSK signature and tumor fitness signature scoring were both performed using the “sc.tl.score_genes” function in SCANPY^38^, which was computed finding a set of 100 background genes in the same expression bin (the range of expression of all genes divided by 30) for each gene in the signature. The TSK signature gene set, and tumor fitness gene set was derived from previous analysis, respectively^39,40^.

### EMT-score calculation

The epithelial-mesenchymal transition (EMT) score was calculated using the Single Sample Gene Set Enrichment Analysis (ssGSEA) method from the gseapy^41^ (v1.1.4) package. The analysis utilized the HALLMARK_EPITHELIAL_MESENCHYMAL_TRANSITION gene set from the Molecular Signatures Database (MSigDB)^42^.

### Designation of clonal regions for mutation rate estimation

The Ct (tumor) region of cSCC-1 was initially defined using SpaGCN (v1.2.7), based on gene expression and spatial coordinates. Spots within likely tumor clusters (TSK score > 0.2) were selected as potential Ct-region spots. To smooth the Ct-region boundaries, we used a k-dimensional tree (k-d tree)^43^ approach using the cKDTree method from scipy.spatial (v1.13.1) to identify the six nearest neighbors for each spot. A spot was then classified as Ct if at least two of its neighbors were labeled as tumor. To define regions for the non-tumor Cp and Cn clones, we first identified the normal epithelial region using SpaGCN. Founder mutations specific to Cp and Cn were then used to label the seed spots for each clone within the identified normal epithelium. The regions were subsequently refined and expanded using a k-nearest neighbors’ algorithm^44^ with 6 neighbors, excluding the spot itself, implemented via the NearestNeighbors module from sklearn.neighbors (v1.4.1) with default parameters.

### Alignment of spots across adjacent tissue slices

Three-dimensional (3D) tissue reconstruction was performed using STAligner^45^ (v1.0.0) to integrate consecutive sections from the same donor. STAligner identifies mutual nearest neighbor (MNN) correspondences between spots from adjacent slices with similar gene expression profiles in the landmark domain, and uses these correspondences to guide 3D spatial registration via the Iterative Closest Point (ICP) algorithm. This analysis was applied to 10x Genomics Visium datasets, including skin cSCC samples (Fig. 4) and human DLPFC samples (Fig. 5).

### Differential expressed genes (DEG) analysis

DEG analysis was performed using the two-tailed Wilcoxon rank-sum test implemented in the rank_genes_groups() function of the python SCANPY^46^ (v1.9.8) library, where genes were filtered to include only those expressed in at least three cells. The log2 fold change (log2FC) was calculated as the difference in mean expression between the tumor cell-of-origin clone (Ci) and the normal epithelial clone (Cn). We identified DEGs using a cutoff of an adjusted p-value of 0.05 and a log2 fold change threshold of 0.1. For 12 DLPFC slices, we first removed the spatial information from each slice and used Harmony^47^ to eliminate batch effects and merge these slices. Subsequently, we identified DEGs between less-proliferative and more-proliferative clones using the wilcox_limma mode in Seurat’s FindMarkers function (v5.2.0)^48^, with a significance threshold of p-value < 0.05 and |FC|>1.5.

### Phylogenetic fitness score calculation

The phylogenetic fitness scores^49^ were computed using the local branching index (LBI)^50^, which incorporates both forward and backward passes across the phylogenetic tree with a time scale parameter for exponential decay. The resulting fitness scores were assigned to each spot, reflecting its position within the phylogenetic tree based on mutation patterns and quantifying the evolutionary success or adaptiveness of a given spot within the tissue slice.

### Tumor phylogenetic fitness–associated differential expression

Genes whose expression varied along the fitness continuum were identified using nested linear regression models. For each tumor and each gene j detected in more than 10 cells, we first normalized raw UMI counts by the cell-level library size and computed log-transformed expression as log(1+e_i,j_)where e_i,j_ denotes the normalized expression of gene j in cell i (or spot i). To account for technical variation in sequencing depth and complexity, we included the per-cell number of detected genes as a size factor size_factor_i_. We then fitted two Gaussian linear models:

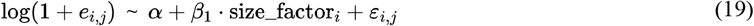

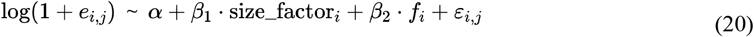

Where f_i_ is the fitness score of cell_i_. A likelihood ratio test (LRT, 1 degree of freedom) was used to compare the full model to the reduced model and to assess whether inclusion of the fitness term β_2_ significantly improved model fit for each gene. Log2 fold-changes were additionally computed by comparing the mean normalized expression of each gene between cells in the top and bottom quantiles of the fitness score distribution.

### Cell-cell communication analysis

To analyze cell–cell communication in Fig. 4, we applied CellChat^51,52^ (v2.1.2) to spatial transcriptomics data processed with Spotiphy (v0.3.1)^36^. For each sample, tumor keratinocytes were stratified into high- and low-fitness spots based on a phylogeny-derived fitness score discussed above, using the median value as the threshold. We then used DBSCAN^53^(v1.2.4) with K-nearest neighbors (KNN) to identify spatially coherent clusters of high-fitness tumor regions (associated with greater clonal expansion) and low-fitness regions.

Parameters of DBSCAN were optimized over a grid: neighborhood radius (ε from 30 to 60, step 5) and minimum points (minPts from 5 to 12). The combination yielding the optimal trade-off between cluster separation and noise was selected. For each group, the cluster containing the largest number of spots was retained. These two spatially defined clusters—high-fitness (associated with greater clonal expansion) and low-fitness—served as input groups for CellChat. We inferred ligand–receptor-mediated communication networks from normalized expression data and cell-type annotations. All detected interactions and their strengths were retained for comparative analysis between the high- and low-fitness groups.

### Density plot of tumor fitness scores

Spatial coordinates, cell-type labels, and expression values were generated with Spotiphy^36^, as discussed previously. We analyzed two continuous metrics: the phylogenetic fitness score based on the phylogeny tree, and the tumor signature fitness score based on gene-expression modules. Analysis was limited to spots located within the tumor region that had valid spatial coordinates. Two-dimensional smoothed density maps were generated for each score using seaborn.kdeplot^54^ (v0.13.2). Kernel density estimation (KDE) bandwidth was determined automatically using Scott’s rule (the default in seaborn.kdeplot, implemented via SciPy^55^ Gaussian KDE), which adapts to the underlying data distribution. Local neighborhood information was derived using the NearestNeighbors function from scikit-learn^56^ (v1.7) to smooth the spatial density mapping.

## Extended Data Figures

**Extended Data Figure 1.**
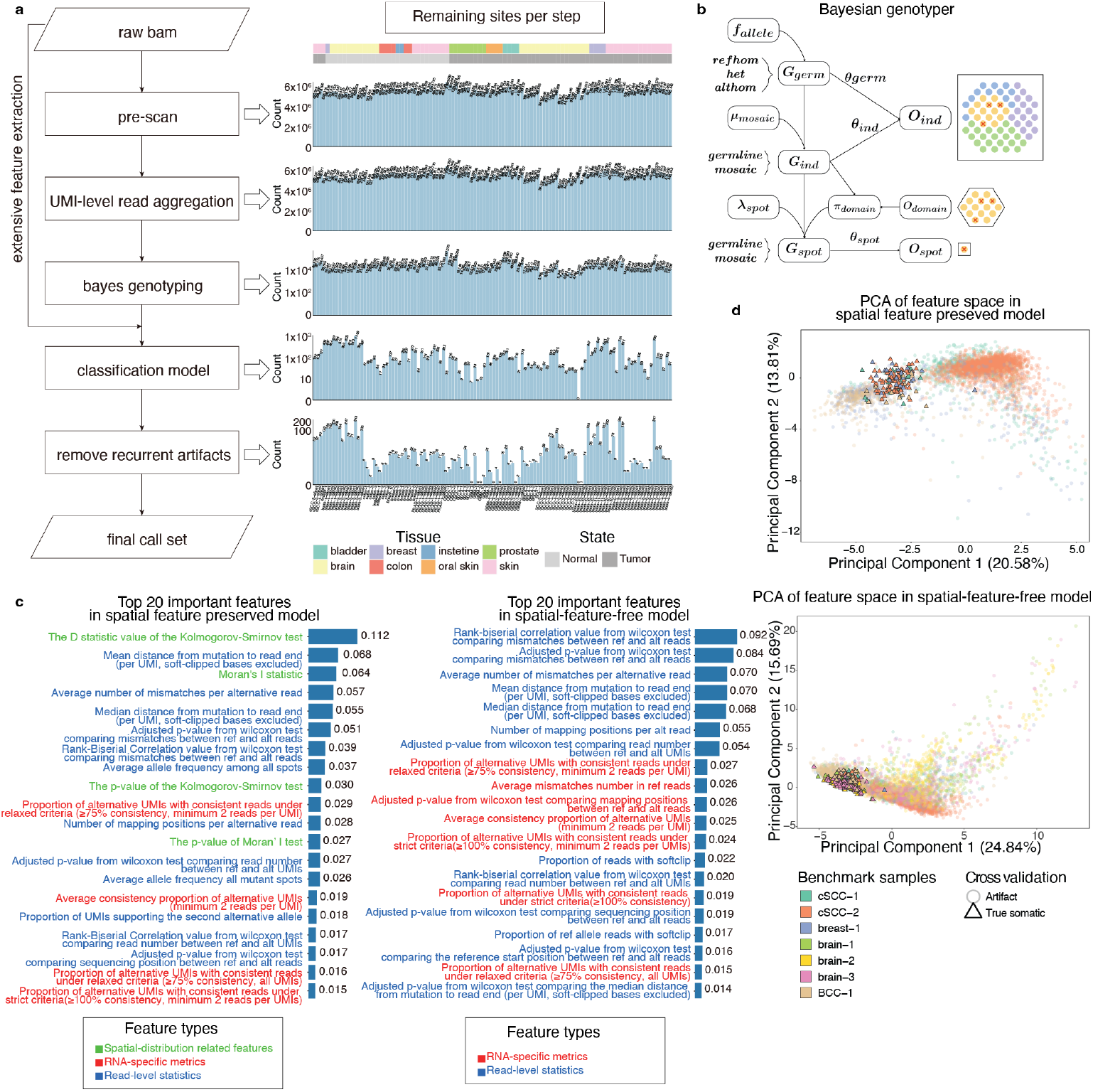
Overall design of SpaceTracer. (**a**) Stepwise design and mutation retention in SpaceTracer across multiple samples analyzed. (**b**) Hierarchical Bayesian Framework of the genotyper. (**c**) Top 20 important features of the Random Forest models for spatial feature preseved model and spatial feature free model. (**d**) PCA projection of the top 20 features with the highest feature importances, to distinguish somatic variants from artifacts. SpaceTracer robustly distinguishes somatic mutations from sequencing artifacts in the benchmark datasets, with or without spatial features. The data sources are described in detail in Supplementary Table 2.

**Extended Data Figure 2.**
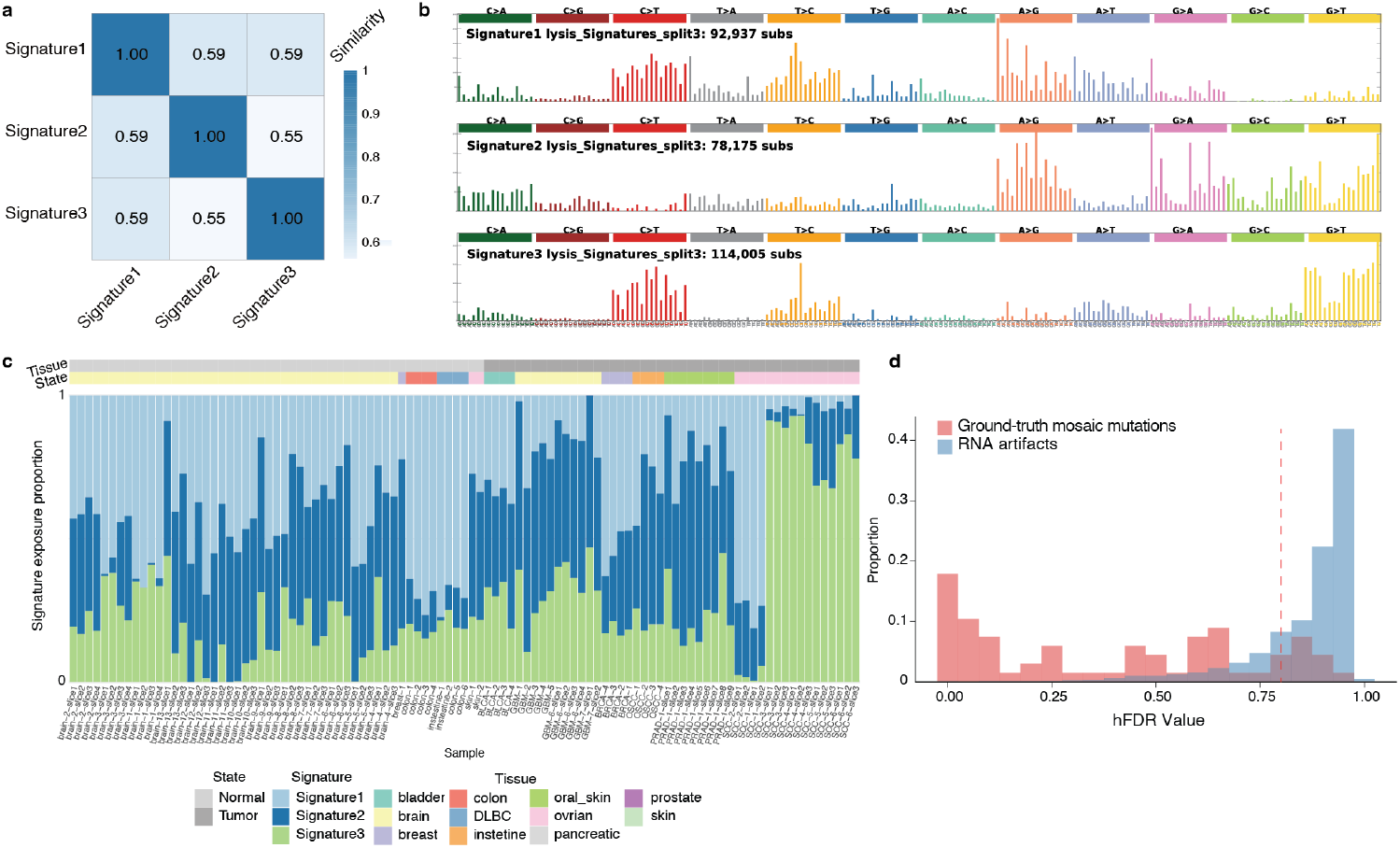
Artifact mutational signature deconstruction and filtration. (**a-b**) Artifact mutational signatures were extracted from 107 slices of spatial transcriptomics data from 50 independent individual samples. Pairwise cosine similarity of NMF-derived artifact signatures was shown in (**a**) and 192-category mutational profiles of the three artifact signatures were shown in (**b**). (**c**) Artifact signature exposure across samples. (**d**) Distribution of hFDR values for ground-truth lysis artifacts versus validated somatic mutations. The dashed vertical line at x= 0.8 indicates the threshold used for hFDR filtering.

**Extended Data Figure 3.**
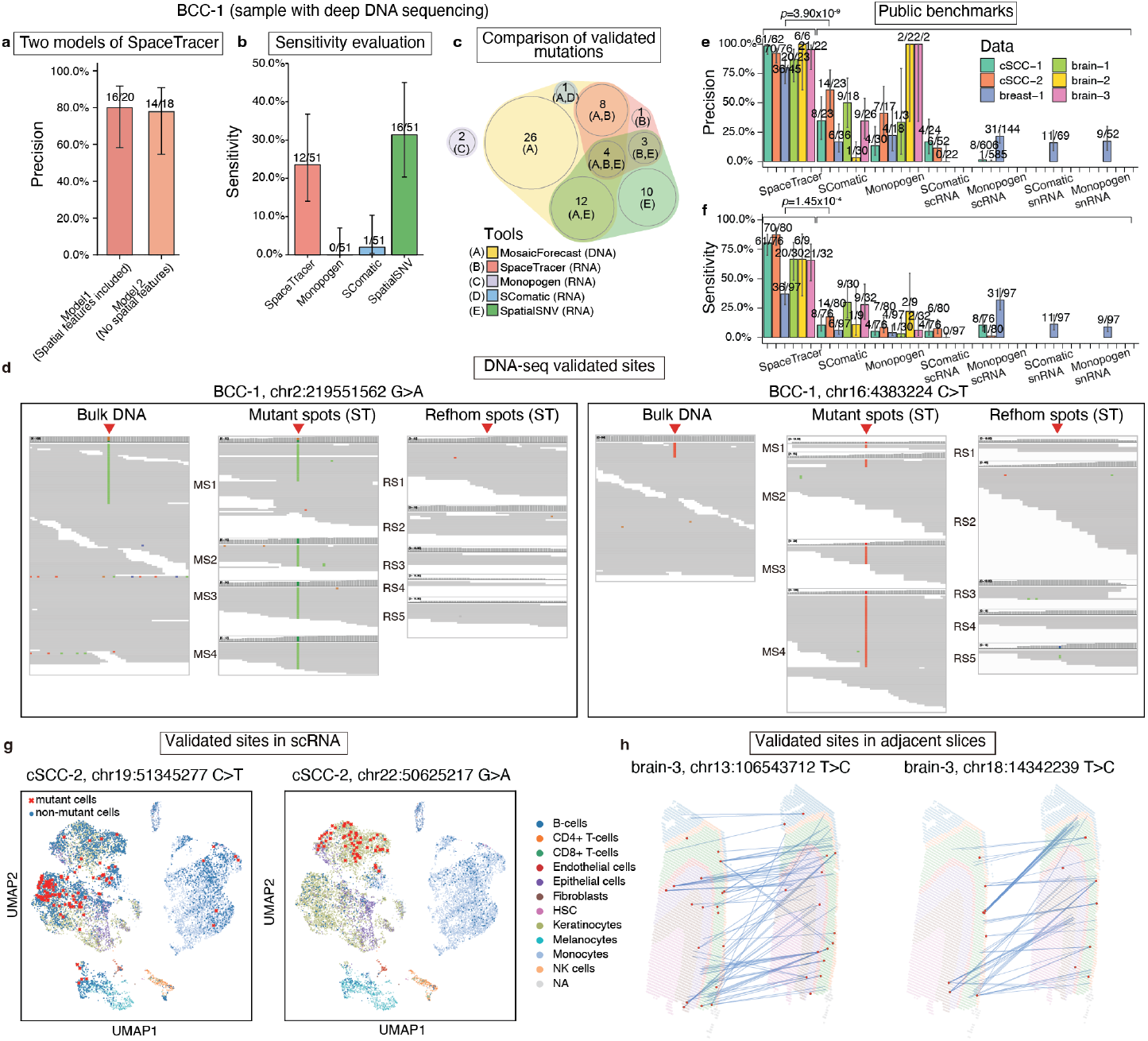
Benchmarking of mutation detection. (**a**) Two random forest (RF) models of SpaceTracer, trained with and without spatial related features, achieved comparable accuracy. (**b**) Sensitivity of each tool is benchmarked against a “ground-truth” set of mutations identified from the two 200× bulk WGS data of two adjacent tissue slices. See Methods for more details. (**c**) Venn diagram comparing validated mutations detected by different software tools. Notably, in addition to mutations detected directly in DNA, multiple mutations were identified from RNA sequencing data and subsequently validated by DNA data, highlighting the unique advantage of in-situ detection of mutations directly RNA-seq. (**d**) IGV plots of validated somatic SNVs in matched deep DNA and spatial transcriptomics data. (**e-f**) Validation and recall rates across public benchmarks. P-values were calculated with one-tailed t-test. Error bars represent 95% CIs calculated with binomial sampling. Values above bars indicate performance metrics. In the precision panel (**e**), the fraction represents validated calls out of total calls made by the tool. In the sensitivity panel (**f**), the fraction represents validated calls detected by the tool out of the total true mutations. The precision and recall comparisons of public benchmarks did not include SpatialSNV because it typically generates tens of thousands of mutations per sample, making manual read-quality validation—required for cross-validation in public datasets—impracticable here. (**g**) Illustrative somatic SNVs validated using paired RNA-seq data. The mutant cells are marked by red Xs. (**h**) Illustrative somatic SNVs validated using adjacent slices of spatial transcriptomics data from a human brain cortex sample with two adjacent slices. Red dots represent mutant spots, and the connecting lines indicate the nearest spots between the two slices, calculated with STAligner^57^ based on both location information and expression data. The mutation is present in both slices at similar locations and exhibits a similar gene expression profile.

**Extended Data Figure 4.**
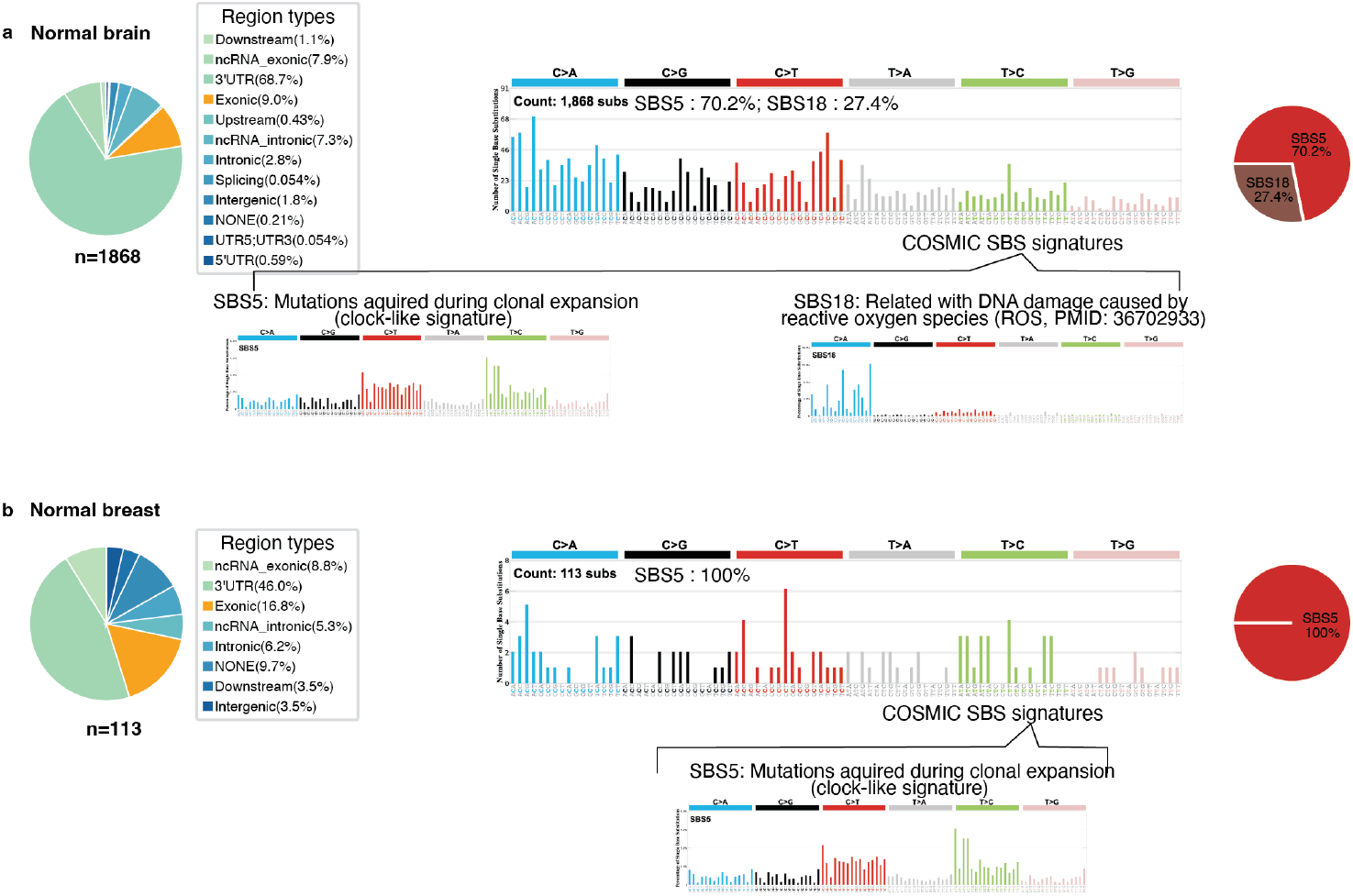
Functional annotation and Mutation signature analysis across benchmark tissues. After filtering likely artifactual mutations, the remaining signatures align with known biological processes and published mutation signatures from DNA sequencing, illustrated by normal brain (**a**), and normal breast **(b**).

**Extended Data Figure 5.**
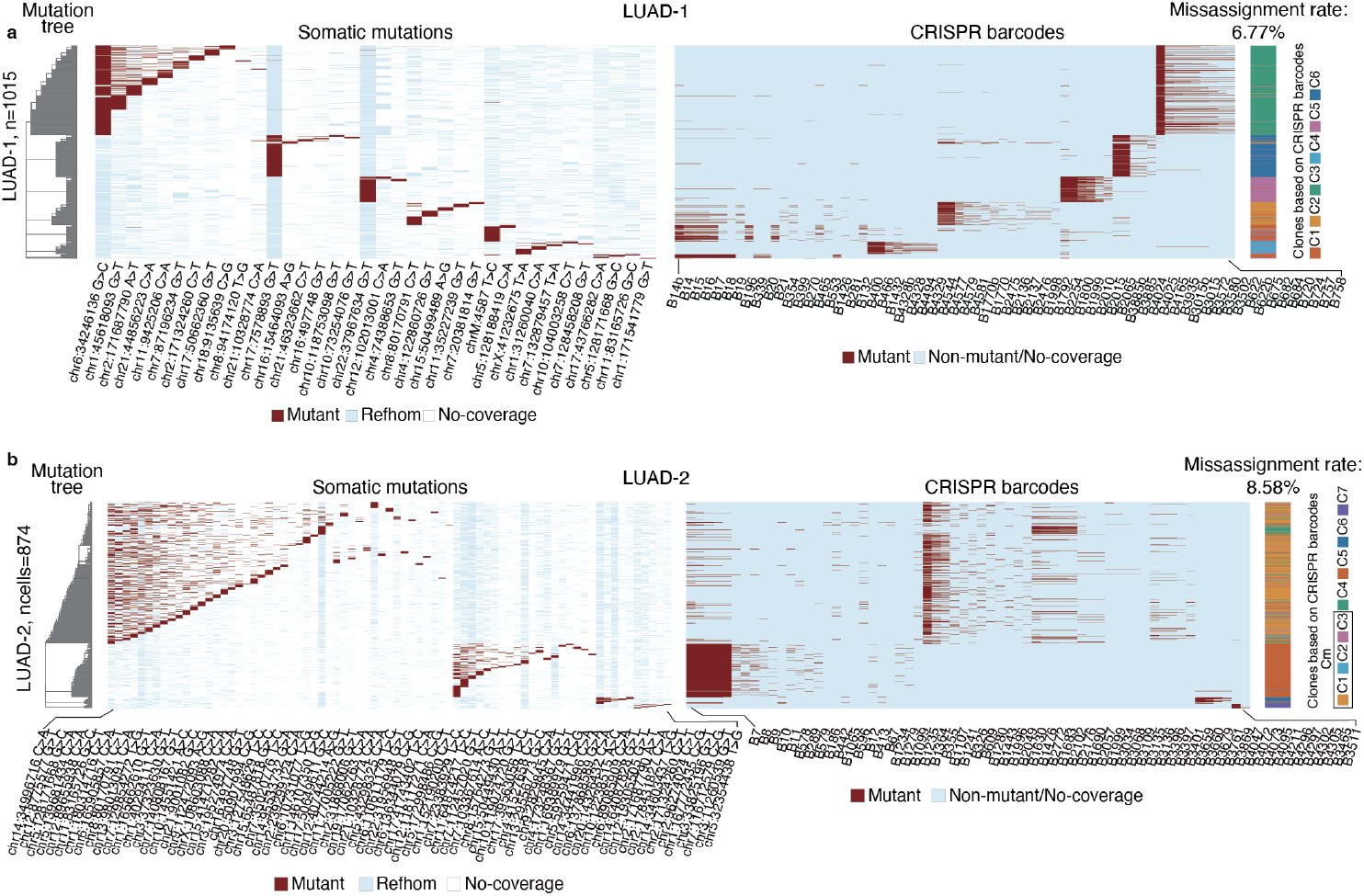
Benchmarking of phylogeny reconstruction. (**a**) Phylogenetic concordance for LUAD-1 between somatic SNV derived and CRISPR barcode (ground truth) lineages. This is an expanded view of Fig. 3b, presenting all annotated somatic mutations and barcodes. (**b**) Phylogenetic concordance for LUAD-2 between somatic SNV derived and CRISPR barcode (ground truth) lineages. This is an expanded view of Fig. 3d, presenting all annotated somatic mutations and barcodes.

**Extended Data Figure 6.**
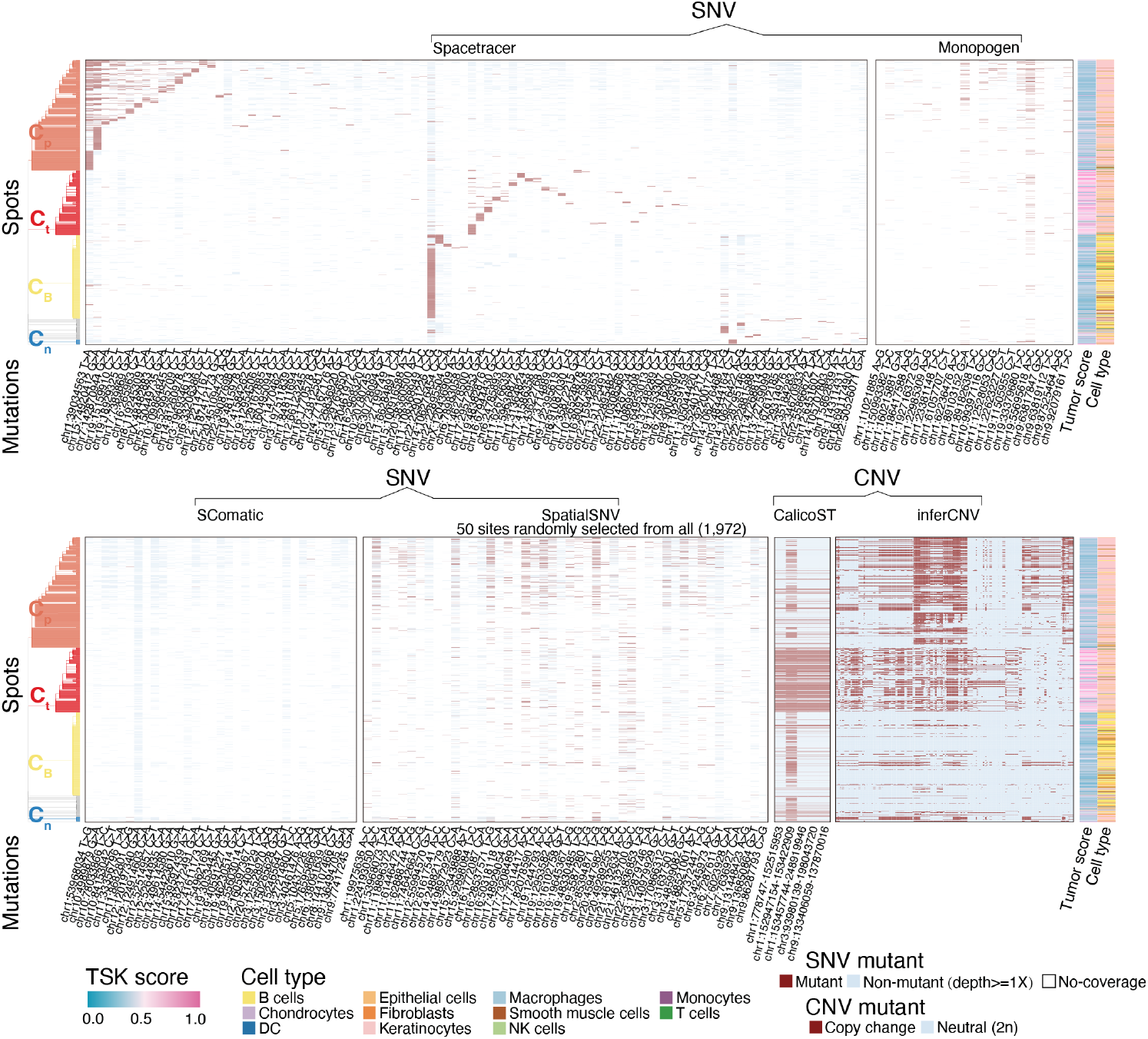
Somatic SNVs and CNVs detected by different software tools from cSCC-1. Each row in the heatmap represent one cell and each column represent one mutation. For visual clarity, 50 mutation sites were randomly selected from thousands of calls generated by spatialSNV. SpaceTracer generate mutation calls of clean clonal structures concordant with CNV calls generated by calicoST. Somatic SNVs called by alternative software tools are limited by either high noise or low suitability for lineage tracing.

**Extended Data Figure 7.**
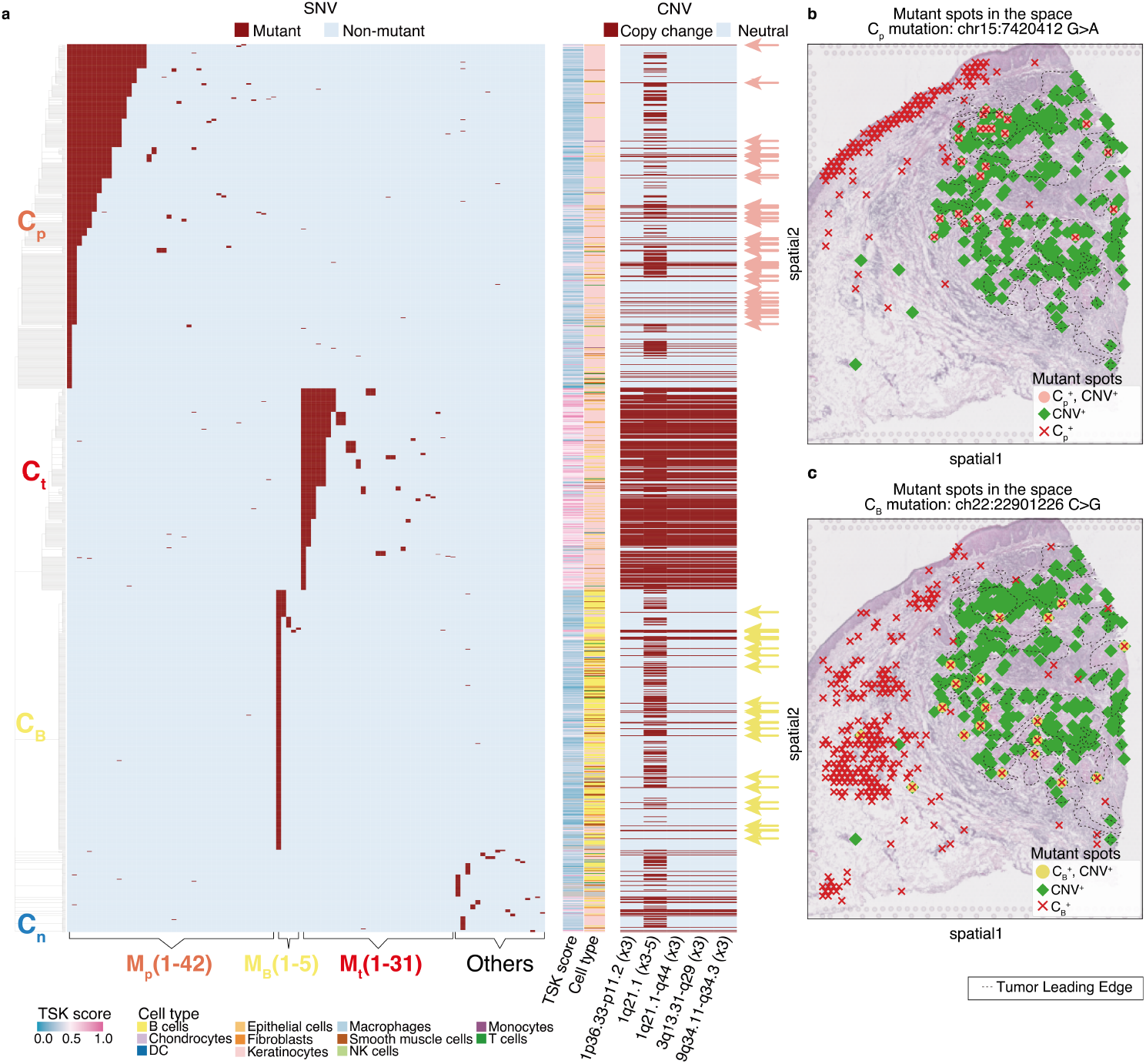
Lineage discrepancies between CNVs and SNVs result from the multi-cellular resolution of spatial sequencing. Spatial profiling captures distinct cell lineages within the same spot, explaining the co-detection of cancer CNVs and lineage-specific single-nucleotide variants (SNVs). **(a)** Heatmap of mutant spots (rows) and somatic mutations (columns). CNVs detected with CalicoST are shown to the right. Arrows highlight spots containing both cancer CNVs and non-tumor-lineage SNVs: Cp mutations from non-tumor epithelium (pink arrows) and CB mutations from B cells (yellow arrows). **(b)** Spatial distribution of Cp-founder mutant spots and cancer-CNV mutations. Spots co-carrying the C_p_ founder mutation and cancer CNVs localize within the tumor region, suggesting migration of C_p_-lineage cells into the tumor and subsequent co-capture with tumor cells. **(c)** Analogous to panel **b**, showing migration of B cells into the tumor region and co-capture of C_B_-mutant cells with CNV-positive tumor cells.

**Extended Data Figure 8.**
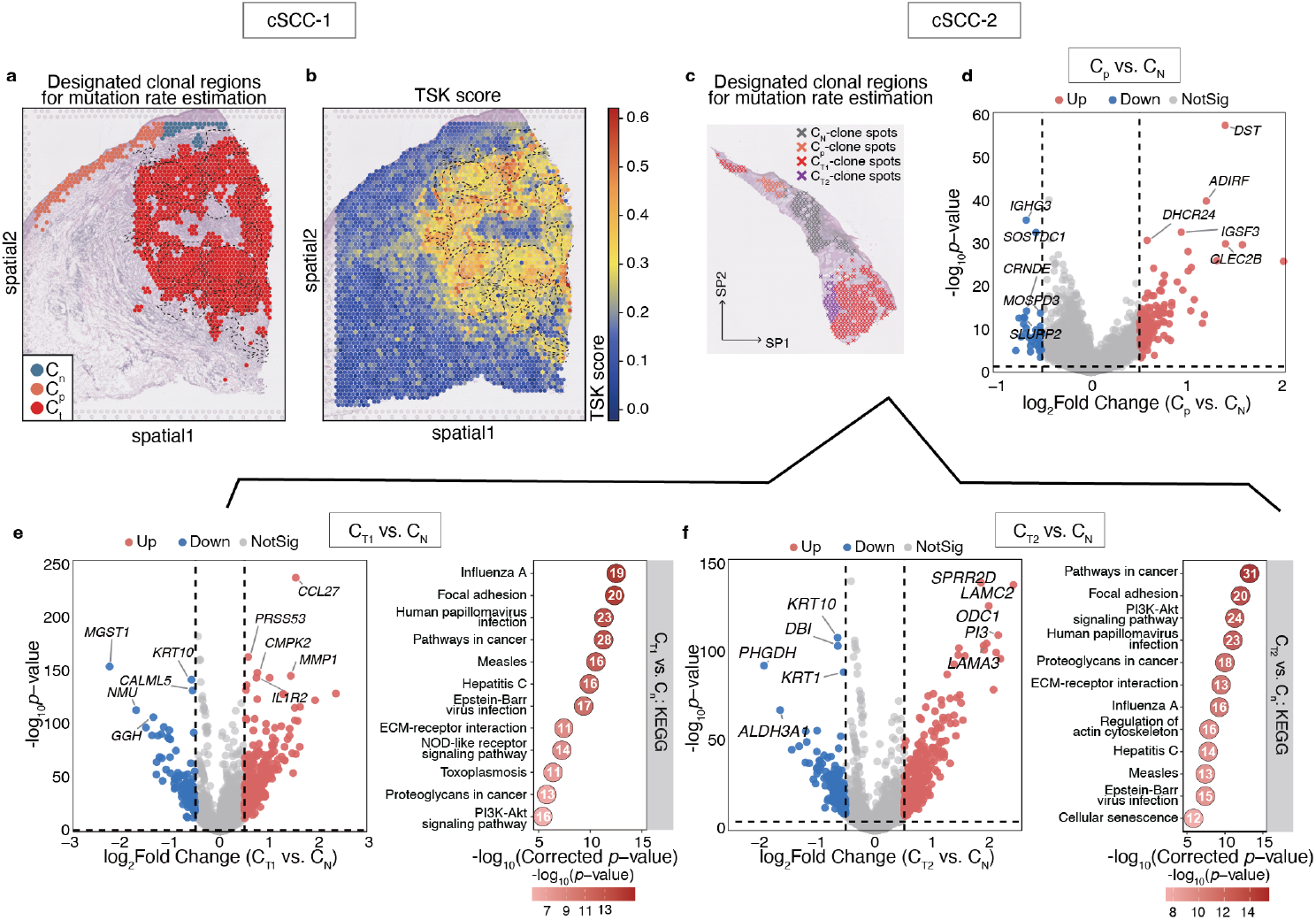
Spatial distribution of tumor and normal epithelial clones and lineage-coupled gene expression changes. **(a)** Designated clonal regions of epithelial clones including C_p_, C_n_, and C_t_ of cSCC-1 for mutation rate estimation. See Methods for more details. (**b**) The spatial distribution of tumor TSK scores of cSCC-1. **(c)** Designated clonal regions of epithelial clones including C_p_, C_N_, C_T1_ and C_T2_ of cSCC-2 for mutation rate estimation. See Methods for more details. (**d**) Differential gene expression analysis of C_p_ vs C_N_ in cSCC-2. (**e-f**) Volcano plots showing differential expressed genes and pathways, by comparing C_T1_ vs. C_N_ and C_T2_ vs C_N_.

**Extended Data Figure 9.**
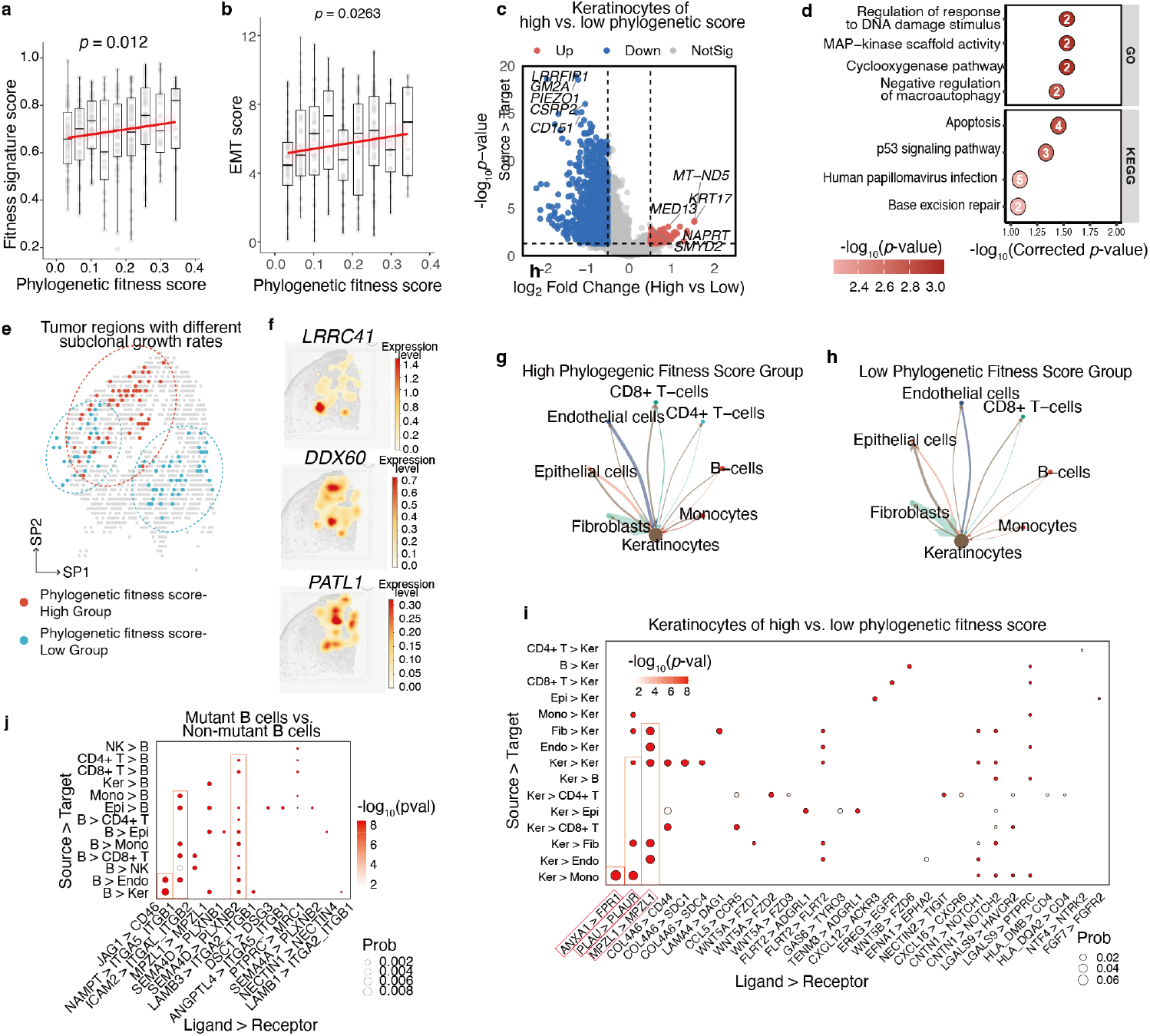
Decoding tumor progression and the tumor-immune ecosystem. (**a**) Linear regression analysis reveals a significant positive correlation between the expression-based tumor fitness signature score and the phylogenetic fitness score (p-value was derived from the Spearman correlation analysis). (**b**) Linear regression analysis reveals that tumor keratinocytes with higher subclonal growth rates (higher phylogenetic fitness score) obtain significantly higher tumor Tumor Epithelial-Mesenchymal Transition (EMT) score (p-value was based on a two-sided t-test of the regression slope). (**c**) Volcano plot of differentially expressed genes in tumor keratinocytes with high versus low phylogenetic fitness. Top significantly up- and down-regulated genes are highlighted. (**d**) Top pathways enriched in tumor keratinocytes with high phylogenetic fitness. (**e**) Tumor regions of high and low phylogenetic tumor fitness were identified using a K-nearest neighbors (KNN) approach (see Methods for details). (**f**) Three top-ranked genes whose expression levels were correlated with phylogenetic fitness scores. The associations were measured with linear regression (see Methods). (**g-h**) Cell–cell communication network of tumor keratinocytes. Arrow width indicates interaction strength. Arrows point from ligand-expressing (source) cells to receptor-expressing (target) cells. (**i**) ligand-receptor pairs significantly enriched in tumor keratinocytes of high versus low fitness scores are highlighted here. (**j**) Analysis of differential gene expressions of mutant B cells versus non-mutant B cells.

**Extended Data Figure 10.**
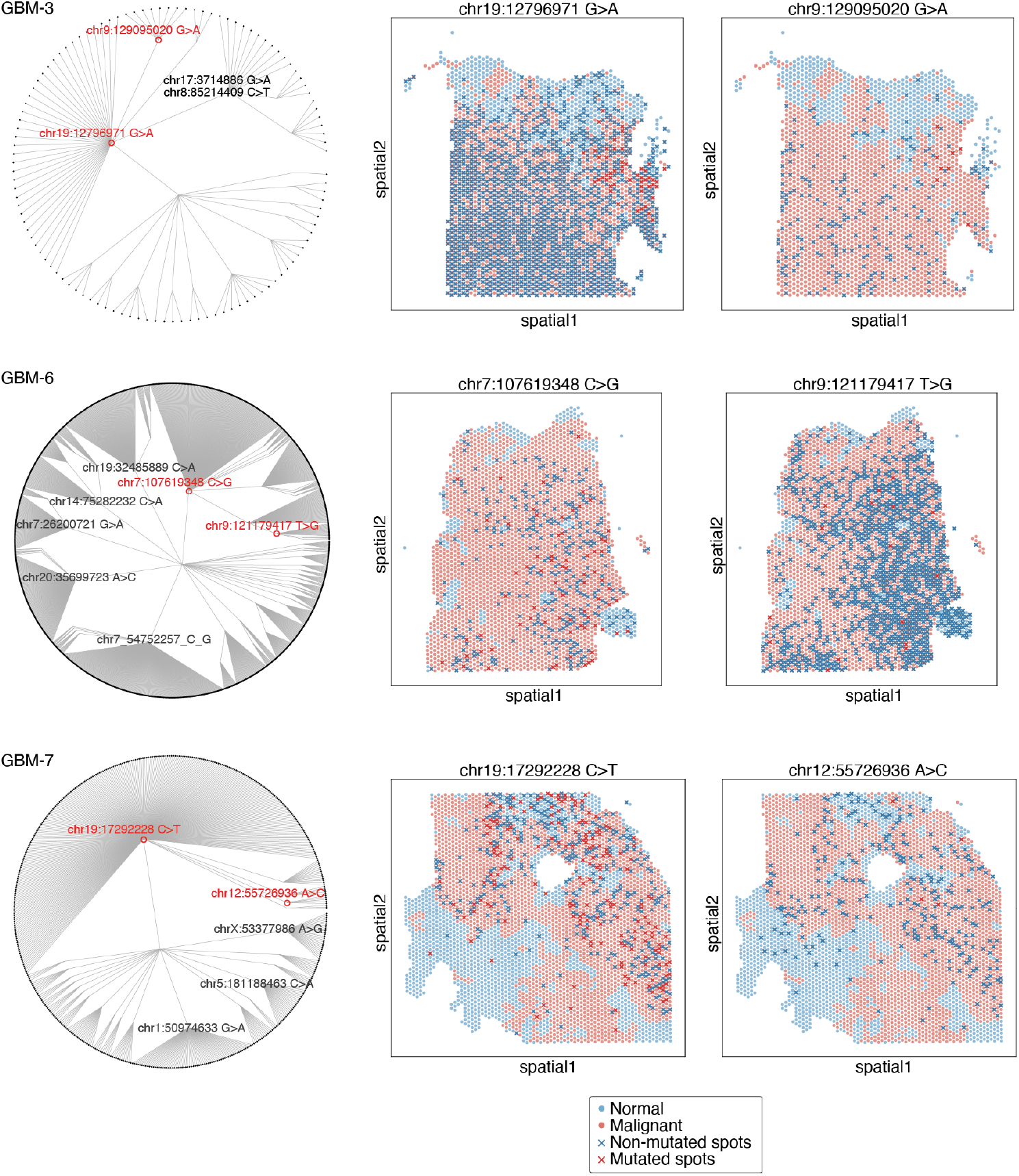
SpaceTracer delineates tumor initiation and progression in glioblastoma samples. (Left) Circular phylogenetic trees reconstructed from three independent glioblastoma samples. Only a subset of mutations is shown from each sample, for visual clarity. (Right) Spatial mapping of two somatic mutations from the same tumor lineage are shown in each row; including an earlier mutation (middle panel) and a later, progression-associated mutation (right panel). Data recourses are listed in Supplementary Table 2.

**Extended Data Figure 11.**
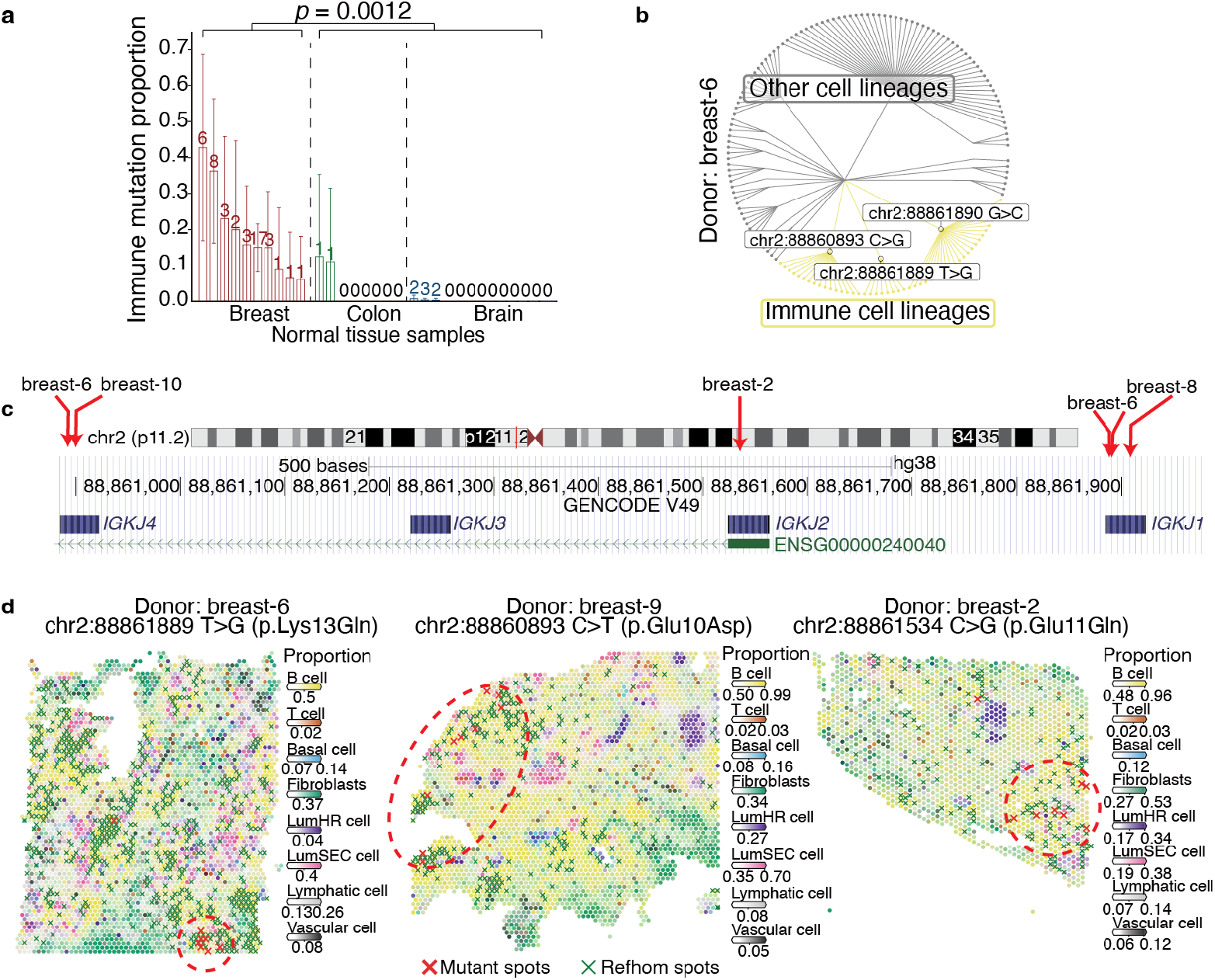
Tissue-reside immune cells across normal tissue samples. (**a**) Among the three types of normal human tissues analyzed, breast tissues harbor a significantly higher proportion of immune cell mutations (two-tailed t-test). The number of immune cell mutations detected in each sample is annotated on the bar. (**b**) Circular phylogeny tree reconstructed with mutations from a normal breast sample, P67. (**c**) Spatial clustering of BCR-mutant cells. Cells harboring somatic mutations within the variable regions of BCR exhibit significant spatial clustering in normal breast tissue.

